# OncoProExp: An Interactive Shiny Web Application for Comprehensive Cancer Proteomics and Phosphoproteomics Analysis

**DOI:** 10.1101/2025.03.06.641407

**Authors:** Edris Sharif Rahmani, Prakash Lingasamy, Soheyla Khojand, Ankita Lawarde, Sergio Vela Moreno, Andres Salumets, Vijayachitra Modhukur

## Abstract

Cancer research has been revolutionized by mass spectrometry (MS)-based proteomics, enabling large-scale profiling of proteins and post-translational modifications (PTMs) to identify critical alterations in cancer signaling pathways. However, the lack of comprehensive, userfriendly platforms for integrative analysis limits efficient data exploration, biomarker identification, and translational insights. To address this gap, we developed OncoProExp, a Shiny-based interactive web application designed for in-depth exploration of cancer proteomes and phosphoproteomes. OncoProExp offers robust workflows for data preprocessing, including missing value imputation and statistical filtering. The platform features interactive visualizations such as principal component analysis (PCA), hierarchical clustering heatmaps, and gene set enrichment analysis (GSEA), enabling detailed functional annotation. Differential expression analysis to identify differentially expressed proteins (DEPs) and phosphoproteins (DEPPs) facilitating the discovery of potential biomarkers and therapeutic targets. The application supports survival analysis and pan-cancer exploration using clinical and proteome/phosphoproteomic datasets. OncoProExp incorporates state-of-the-art predictive modeling using machine learning algorithms, including Support Vector Machines (SVMs), Random Forests, and Artificial Neural Networks (ANNs) for cancer risk stratification, achieving near-perfect accuracy in multi-cancer and single-cancer classification. These models are enhanced by SHapley Additive exPlanations (SHAP) for interpretability. To enhance its translational utility, the platform supports user-uploaded data and enables protein-protein interaction analysis, pathway enrichment analysis, cancer drug relevance evaluation, and clinical annotation using curated cancer-specific datasets. OncoProExp is deployable via Docker containers, ensuring flexible and scalable integration into individual servers. Its utility has been demonstrated using Clinical Proteomic Tumor Analysis Consortium (CPTAC) datasets, showcasing its potential to advance cancer biomarker discovery, risk stratification, therapeutic target identification, and personalized treatment strategies. OncoProExp is freely accessible at https://oncopro.cs.ut.ee/ without login requirements, offering a comprehensive resource for translational cancer research.

## 1. Introduction

Cancer is a heterogeneous disease driven by complex genetic, epigenetic, proteomic, and metabolomic alterations that influence tumor initiation, progression, and metastasis [1–3]. Despite advancements in molecular oncology, the intricate landscape of cancer biology remains challenging for accurate diagnosis and effective therapeutic interventions [4–6]. The diversity of molecular alterations across different cancer types leads to variable treatment responses, underscoring the need for sophisticated tools to dissect tumor heterogeneity and support personalized therapies.

Conventional genomic and transcriptomic analyses often fail to capture the dynamic nature of protein activity and post-translational modifications (PTMs), which are critical determinants of cellular function and disease progression. In contrast, proteomics directly quantifies proteins, offering a more accurate representation of functional cellular states and genotype-phenotype relationships [7,8]. RNA expression data, often used as proxies for protein expression, predict protein levels with limited accuracy (around 40%), offering little insight into PTMs and alternative splice variants [9,10]. PTMs, including phosphorylation, acetylation, and ubiquitination, occur post-synthesis and are not encoded by mRNA, significantly influencing protein function and interactions. Moreover, recent studies have found new microproteins encoded by previously classified non-coding regions that act as oncogenic drivers or tumor suppressors [11,12]. Therefore, direct proteomic analysis provides a deeper understanding of tumor biology compared to transcriptomic or genomic approaches [13–16].

Mass spectrometry (MS)-based proteomics and phosphoproteomics enable large-scale profiling of protein expression and PTMs, uncovering critical mechanisms in oncogenic signaling, immune evasion, drug resistance, and metabolism [17–19]. Proteomics characterizes protein abundance, interactions, and modifications, whereas phosphoproteomics specifically examines phosphorylation events that drive tumorigenesis and drug resistance [20,21]. The application of these approaches has identified novel biomarkers and mechanisms of drug resistance, advancing precision oncology [22,23]. Large-scale initiatives, such as the Clinical Proteomic Tumor Analysis Consortium (CPTAC), have generated extensive proteomic datasets. However, analyzing these datasets remains computationally challenging owing to their data complexity, need for specialized bioinformatics expertise, and requirement for advanced computational skills. Consequently, these initiatives are not accessible to the broader research and clinical communities.

AI and machine learning (ML) have emerged as powerful tools for extracting meaningful insights from multi-omics datasets [24]. AI Techniques such as Artificial Neural Networks (ANNs), Support Vector Machines (SVMs), and Random Forests (RFs), have shown improved cancer prediction, prognosis, and patient stratification using proteomic and genomic data [25,26]. Integrating AI with high-throughput technologies and medical imaging has led to FDA-approved diagnostic tools [27–29]. However, challenges remain in data curation, standardization, and development of interpretable AI models validated in clinical settings. Proteomics and phosphoproteomics datasets introduce additional complexity due to missing values, outliers, and non-linear relationships in cancer data [30]. Current bioinformatics platforms facilitate proteomic data exploration, but lack comprehensive workflows that integrate robust preprocessing, differential expression analysis, interactive visualization, AI-driven modeling, and pan-cancer analysis.

To address these challenges, we developed OncoProExp, an interactive Shiny web application designed for comprehensive cancer proteomics and phosphoproteomic analyses in cancer research. OncoProExp offers robust data preprocessing, intuitive visualization, and advanced statistical analysis tailored explicitly for pan-cancer investigations across eight cancer types: CCRCC, COAD, HNSCC, LSCC, LUAD, OV, PDAC, and UCEC. These capabilities align with the growing demand for personalized cancer therapies and precision medicine. The platform ensures high-quality data processing through missing-value imputation, outlier detection, and feature scaling, which are critical for reliable predictive modeling. Users can make use of advanced visualization tools, such as Principal Component Analysis (PCA), Multidimensional Scaling (MDS), and Uniform Manifold Approximation and Projection (UMAP) to explore the relationships between tumor and normal samples. The integration of advanced AI-based predictive models with traditional bioinformatics methods enhances the accuracy of cancer-type predictions, achieving up to 95% accuracy by capturing non-linear patterns in high-dimensional datasets using models such as SVM, RF, and ANN. By combining these features with comprehensive statistical analyses such as Differential Expression Analysis (DEA) and Gene Set Enrichment Analysis (GSEA), OncoProExp provides a comprehensive platform for cancer proteomics and phosphoproteomic analysis. Its user-friendly interface enables researchers to conduct in-depth explorations of cancer-related proteomics data, facilitates biomarker discovery and functional analysis, enhances understanding of cancer biology, and predicts cancer types based on molecular profiles.

## 2. Materials and Methods

### 2.1 Data Collection

Proteomics and phosphoproteomics data were obtained from the CPTAC pan-cancer cohort using LinkedOmics [31] (https://linkedomics.org/login.php#dataSource, accessed May 2024). The datasets included Tandem Mass Tag (TMT)-based quantitative proteomics and phosphoproteomics profiles, with protein and phosphoprotein abundance expressed as log2-transformed ratios relative to a reference sample. This approach allows for precise quantification of relative protein and phosphosite abundance across samples. The inclusion criteria were ≥10 normal samples and ≥60 patient samples per cancer type. Eight cancer types—Clear Cell Renal Cell Carcinoma (CCRCC), Colon Adenocarcinoma (COAD), Head and Neck Squamous Cell Carcinoma (HNSCC), Lung Squamous Cell Carcinoma (LSCC), Lung Adenocarcinoma (LUAD), Ovarian Cancer (OV), Pancreatic Ductal Adenocarcinoma (PDAC), and Uterine Corpus Endometrial Carcinoma (UCEC)—met these criteria. Glioblastoma Multiforme (GBM) and Breast Cancer (BRCA) were also included for survival analysis, despite the lack of normal samples, due to their clinical relevance. All data consisted of log2-transformed MS1 intensities, with paired tumor and normal samples prioritized for direct comparisons. A summary of the datasets, including sample numbers, average age, and proportion of females, is provided in Table 1.

**Table 1:** Overview of datasets used in the current study from CPTAC comprising proteomic and phosphoproteomic datasets.

| Cancer Code | Cancer Type | Average Age | Female (%) | Tumor Samples | Normal Samples |
| --- | --- | --- | --- | --- | --- |
| CCRCC | Clear Cell Renal Cell Carcinoma | 60.79 | 25.24 | 103 | 80 |
| COAD | Colon Adenocarcinoma | 64.43 | 57.8 | 109 | 100 |
| HNSCC | Head and Neck Squamous Cell Carcinoma | 61.44 | 12.96 | 108 | 62 |
| LSCC | Lung Squamous Cell Carcinoma | 65.87 | 20.37 | 108 | 99 |
| LUAD | Lung Adenocarcinoma | 62.6 | 34.55 | 110 | 101 |
| OV | Ovarian Cancer | 58.92 | 100 | 83 | 19 |
| PDAC | Pancreatic Ductal Adenocarcinoma | 64.49 | 43.81 | 105 | 44 |
| UCEC | Uterine Corpus Endometrial Carcinoma | 63.47 | 100 | 95 | 18 |
| GBM | Glioblastoma Multiforme | 57.89 | 44.44 | 99 | None |
| BRCA | Breast Cancer | 59.99 | 100 | 122 | None |

### 2.2 Data Preprocessing

The initial step in data quality control began by eliminating features with substantial missing values, over 50% for the proteome and more than 70% for the phosphoproteome. The Random Forest algorithm, implemented using the missForest package (v1.5) in R [32], was used with default parameters for iterative imputation of the remaining missing values until convergence was attained. Subsequently, the tumor and normal datasets were synchronized with the metadata, retaining only the overlapping features. Gene symbols were standardized using Ensembl database in biomaRt (v2.58.2) [33] to ensure consistency in gene identifiers. In cases where multiple entries corresponded to a single gene, expression values were averaged. The resulting consolidated dataset, comprising both tumor and normal data, was saved as a CSV file for further analysis.

### 2.3 Visualization

For visualization purposes, we focused on the top most 1,000 variable proteins or phosphoproteins based on median absolute deviation (MAD). The resulting plots underwent dimensionality reduction via Principal Component Analysis (PCA), Multidimensional Scaling (MDS), and Uniform Manifold Approximation and Projection (UMAP). Scree plots were used to determine the number of principal components that explained most of the variance. Plots were color-coded by tumor/normal or other metadata categories. Box and density plots revealed value distributions, aiding outlier detection. Heat maps with hierarchical clustering were used to further summarize the expression patterns. When applicable, Gene Set Enrichment Analysis (GSEA) (using fgsea v4.7-12 in R with default parameters) [34] bar plots highlighted significantly enriched pathways among the primary data partitions, offering biological insights into functional processes and pathway activation.

### 2.4 Differential Expression Analysis

Differentially expressed proteins (DEPs) and phosphoproteins (DEPPs) were identified using limma with default parameters (v3.58.1) [35], applying empirical Bayes moderation. Significance thresholds were defined as |log2FC|>0.8 and FDR<0.01. Volcano plots displayed log2FC versus log10 p-values, while mean–standard deviation plots were used to highlight outliers. Box plots and heatmaps summarized DEP/DEPP expression differences between tumor and normal samples.

### 2.5 Enrichment Analysis

Enrichment analysis for DEPs/DEPPs was performed using gprofiler2 v0.2.3, with a set of default parameters [36] to identify functional annotations from Gene Ontology (GO), Kyoto Encyclopedia of Genes and Genomes (KEGG), and Reactome annotations (FDR<0.05). Bar plots were used to visualize pathway significance and gene set sizes. Interactive visualizations were generated using Plotly (v4.10.4) [37], which enables category-specific exploration and visualization. KEGG pathway diagrams were generated using the Pathview package with default parameters to visualize the relationships between DEPs and DEPPs within enriched pathways. Protein-protein interaction (PPI) networks were constructed using string database (STRINGdb v2.14.3, using default parameters) [38] to identify key regulatory proteins and interaction hubs. To evaluate therapeutic relevance, dysregulated proteins were cross-referenced with public drug databases, including DrugBank [39] and the Cancer Drugs Database [40]. Box plots illustrations are provided for the differential expression of drug-targetable genes, highlighting their potential as candidates for therapeutic interventions.

### 2.6 Survival Analysis

Cox proportional hazards models (both univariate and multivariate) were implemented using the default parameters of Survival (v3.7) and survminer (v0.4.9) packages [41,42]. To assess survival differences, patients were stratified into high- and low-expression groups based on the median expression levels of the biomarkers of interest. This median-based stratification ensured balanced group sizes and minimized the bias in the analysis. Survival differences between the high- and low-expression groups were evaluated using log-rank tests. Kaplan-Meier curves were generated to visualize survival probabilities over time and hazard ratios (HR) with 95% confidence intervals (CI) were calculated to quantify the impact of individual biomarkers on survival outcomes. In multivariate analyses, clinical variables such as age, sex, and tumor stage were included as covariates to adjust for potential confounding factors and provide a more comprehensive assessment of the prognostic significance of the biomarkers.

### 2.7 Pan-Cancer Analysis

A comparative analysis of proteomic and phosphoproteomic datasets from eight cancer types (CCRCC, COAD, HNSCC, LSCC, LUAD, OV, PDAC, and UCEC) was conducted using pan-cancer analysis to identify common and distinct dysregulated features between tumor and normal samples. The study compiled differential expression metrics and p-values to highlight inter-cancer expression patterns. Additionally, related survival analyses were performed to pinpoint biomarkers with prognostic significance across multiple cancer types.

### 2.8 Feature Selection

A two-step feature selection process was performed to identify the most relevant diagnostic biomarkers. First, Random Forest-based importance scores were calculated using the random-Forest package (v4.7-1.2) [43] to capture non-linear relationships between the variables. Thus, features with importance scores above a predefined threshold and high variability (based on MAD) were retained, ensuring consistent variations across samples and improving the model’s capacity to detect meaningful biological signals. Second, t-SNE projections (Rtsne package, v0.17) [44] were used to evaluate the ability of selected features to differentiate sample clusters. The top 250 proteins and 100 phosphoproteins were selected for downstream machine learning modeling.

### 2.9 AI-Based Predictive Models

Three machine learning algorithms—SVM (e1071, v1.7-16) [45], Random Forest (randomForest), and ANN (keras, v2.15.0) [46] were applied to predict cancer type from proteomics/phosphoproteomics data. SVM used a linear kernel with cost=3, 80:20 train-test split, and 10-fold cross-validation. Random Forest comprised 100 trees, also using 80:20 splits and 10-fold CV. ANN had four layers (200, 100, 30, 16 neurons) with ReLU activation, dropout (40%, 30%, 10%), L2 regularization in hidden layers, and an Adam optimizer (learning rate=0.0009). The training ended early if the validation loss did not improve for five epochs. The data were split into 80% training, 10% validation, and 10% testing.

### 2.10 Model Evaluation

Metrics included accuracy, sensitivity, specificity, precision, F1-score, and AUC (MBMethPred [47], v0.1.4.2). The calculations for these metrics are as follows:

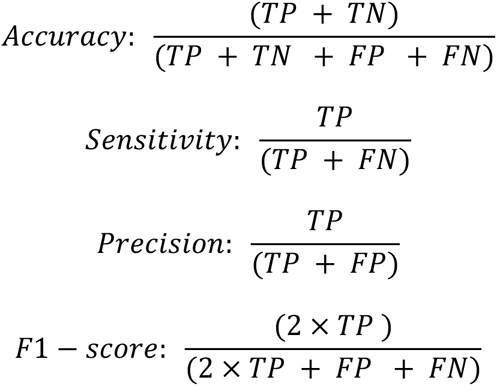

Here, TP stands for true positives, FP for false positives, TN for true negatives, and FN for false negatives, reflecting the correctness of the model’s predictions. Each model was trained and validated using 10-fold CV and an independent test set. This procedure prevents overfitting, and provides robust performance estimates.

### 2.11 Model Interpretation with SHAP

SHapley Additive exPlanations (SHAP) were applied to Random Forest classifiers (scikit-learn, v1.6.0, with default parameters in Python) [48]. TreeExplainer computed the SHAP values per feature, revealing the contribution of each protein or phosphoprotein to sample-level predictions. Mean absolute SHAP values ranked among the top 10 biomarkers, and summary plots showed how high or low expression values nudged classifications, enhancing the interpretability of the AI-based models.

### 2.12 Shiny-based Web Server Development

The interactive dashboard and database were implemented using R Shiny [49] and deployed within a Docker container on a cloud-based hosting platform. This arrangement enables users to easily configure a set-up and seamless integration. Secure communication between the client browser and server is established using the SSL/TLS protocol, ensuring encryption and authentication via HTTPS.

## 3 Results

### Overview and Construction of OncoProExp

OncoProExp is a comprehensive, open-source bioinformatic platform for analyzing cancer proteomics and phosphoproteomic data. This versatile tool is freely available as a web server for quick analysis of small datasets and can also be deployed locally using a Docker container or Shiny-based R package to ensure adaptability, scalability, and reproducibility. Developed using robust R scripts and incorporating various Bioconductor packages, OncoProExp offers a user-friendly experience for researchers of all computational skill levels. The platform features a user-friendly interface with interactive dashboards for data uploading and preprocessing. To improve accessibility, OncoProExp provides downloadable CPTAC datasets for reproducibility, and detailed video tutorials for guidance. Accessible at https://oncopro.cs.ut.ee without requiring login credentials, OncoProExp enables researchers to identify biomarkers, potential therapeutic targets, and molecular insights from proteomics data, facilitating the translation of complex datasets into actionable biological knowledge.

The OncoProExp interface is comprises of four primary components as shown in Figure 1. (1) Data Processing, (2) Differential Analysis, (3) Pan-Cancer and Survival Analysis, and (4) Machine Learning Analysis. These components have been engineered to facilitate seamless exploration of cancer proteomes and phosphoproteomes, enabling researchers to identify biomarkers, potential therapeutic targets, and prognostic indicators. The subsequent sections provide a comprehensive examination of each of these aforementioned components.

**Figure 1.**
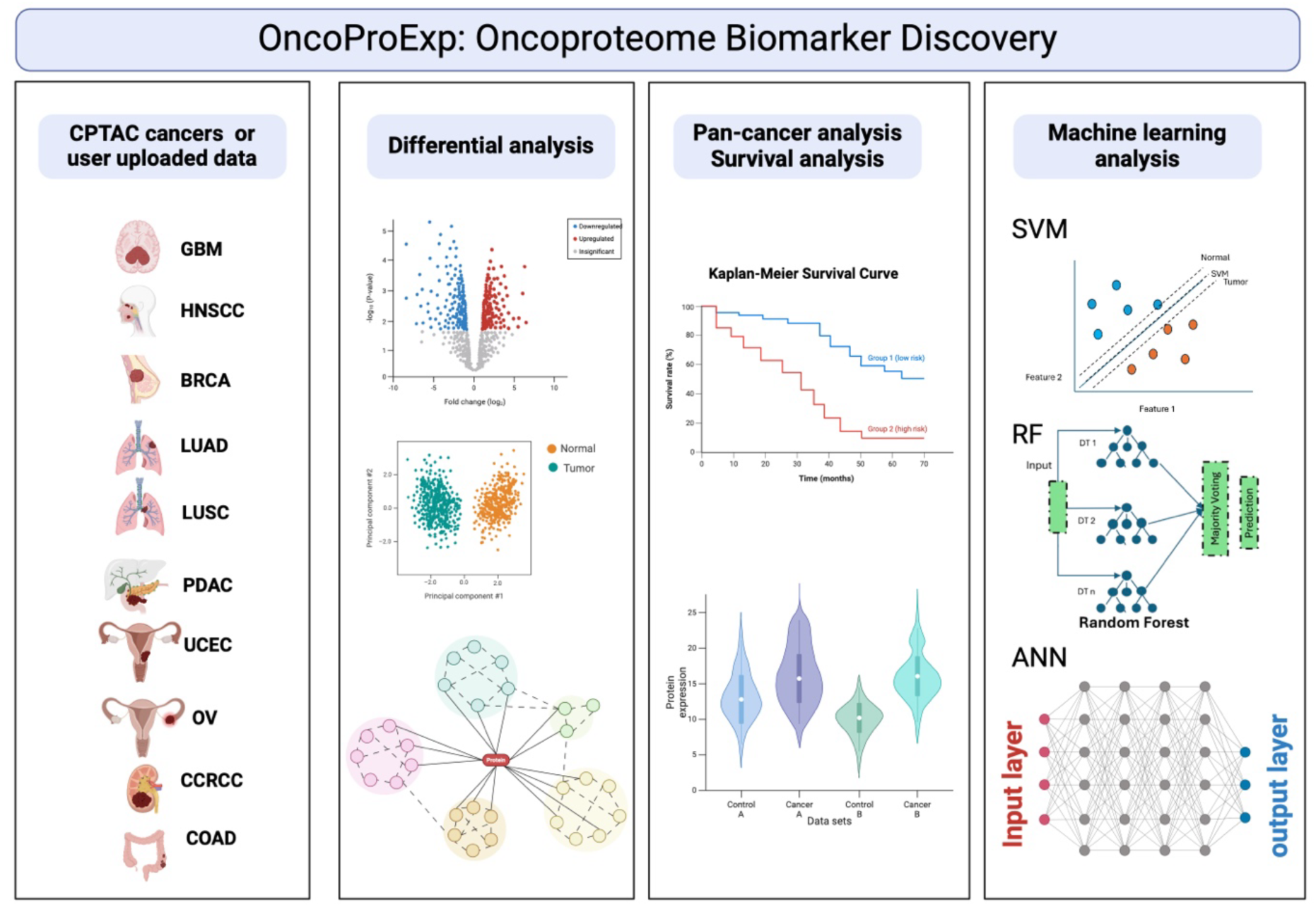
Overview of the OncoProExp platform for cancer proteomics and phosphoproteomic analyses OncoPro-Exp consists of four primary modules: (1) Data Input Module—integrates CPTAC cancer datasets or user-uploaded data, covering multiple cancer types such as GBM, HNSCC, BRCA, LUAD, LUSC, PDAC, UCEC, OV, CCRCC, and COAD. (2) Differential Analysis Module—detects protein expression differences between normal and tumor samples, providing visualizations such as volcano plots, PCA scatter plots, heatmaps, and protein interaction networks, alongside enrichment analyses, KEGG pathway mapping, and drug relevance evaluation using DrugBank and Cancer Drugs Database. (3) Pan-Cancer and Survival Analysis Module—assesses protein expression across various cancer types and its clinical relevance using Kaplan-Meier survival curves and violin plots. (4) Machine Learning Module—applies SVM, Random Forest, and ANN algorithms for biomarker discovery and predictive modeling, facilitating robust and accurate data-driven insights.

### 3.1 Data Processing

OncoProExp enables users to efficiently preprocess proteome and phosphoproteome data by accepting uploaded data in a standardized format or CPTAC data. Users may provide their data as a tab-delimited text file or CSV file, with rows representing genes/proteins or phosphorylation sites, and columns representing samples. The data should consist of log2-transformed ratios or intensities, which quantify the relative abundance of proteins or phosphoproteins compared to a reference sample. For example, a log2 ratio of 1 indicates a 2-fold increase, whereas a log2 ratio of -1 indicates a 2-fold decrease in abundance. Missing values should be marked as NA or left blank. The first column should contain gene symbols (e.g., TP53) or Ensembl IDs (e.g., ENSG00000141510), followed by sample-specific abundance values (Supplementary file 1). After uploading, OncoProExp applies filters to remove samples with high proportions of missing values, retaining all tumor and normal samples from the proteome dataset while excluding specific samples from the phosphoproteome dataset (e.g., one normal sample from OV, eight tumor and nine normal samples from LUAD, six tumor samples from UCEC, and five tumor and four normal samples from LSCC). Gene identifiers are standardized by converting Ensembl IDs to gene symbols and averaging duplicates, resulting in 11,000 unique gene symbols for both datasets. The remaining missing values were imputed using the Random Forest algorithm to ensure robust downstream analysis. The preprocessed data is then saved as a CSV file, ready for further exploration and analysis.

### 3.2 Data Visualization

OncoProExp provides powerful visualization methods to explore differences in proteomic and phosphoproteomic profiles between tumor and normal samples. Users can focus on the top most variable proteins (e.g., 1,000 or any preferred number) or phosphoproteins to generate Principal Component Analysis (PCA), Multidimensional Scaling (MDS), and Uniform Manifold Approximation and Projection (UMAP) plots. These visualizations reveal clear distinctions between sample types, with the first principal component accounting for 35.95% of the variation in HNSCC, 33.84% in PDAC, 24.82% in OV, 20.73% in UCEC, and over 50% in other cancer types (Figure 2A-B).

**Figure 2.**
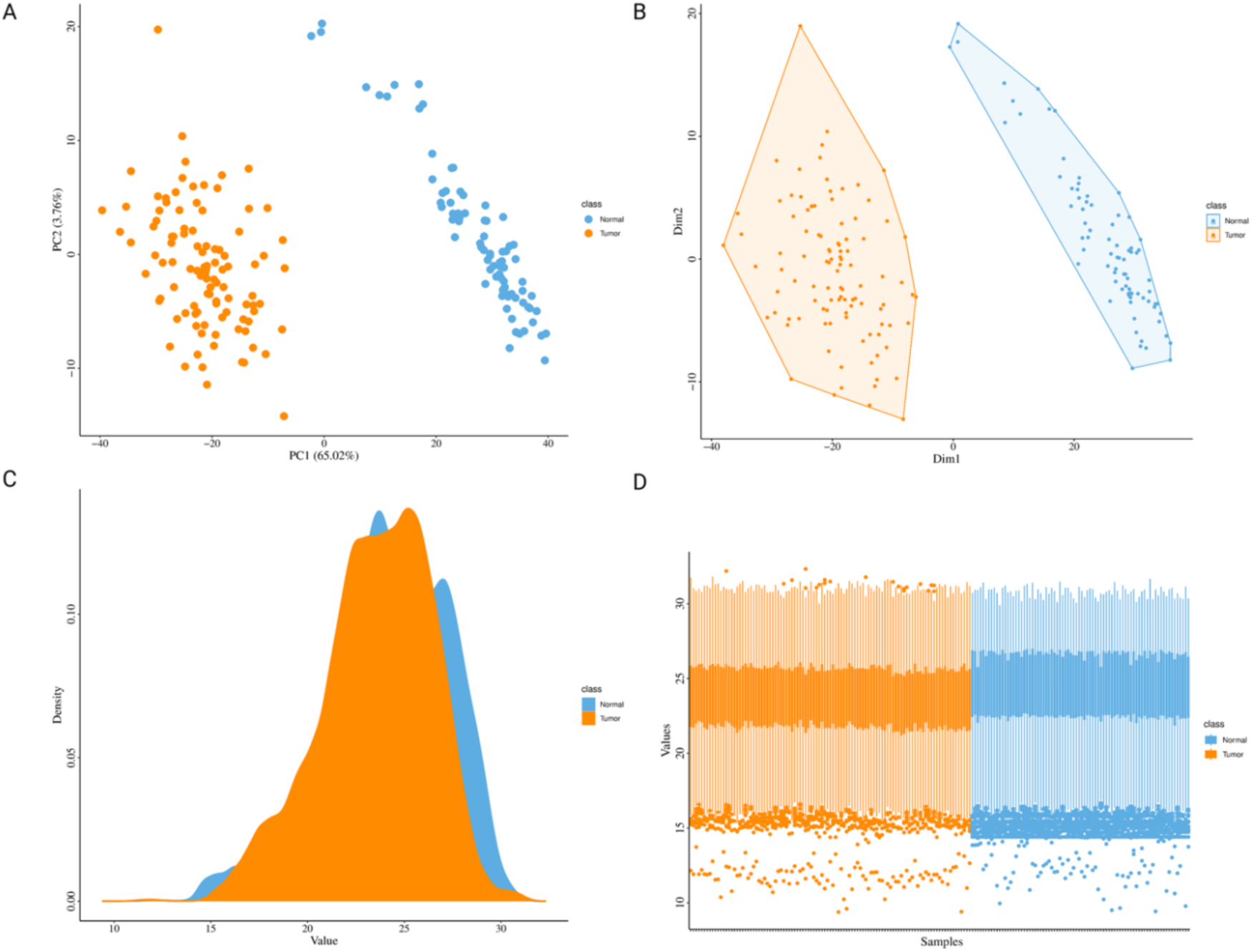
Visualization of CCRCC tumor and normal samples. (A) Principal Component Analysis (PCA) plot showing distinct separation between tumor and normal samples along the first two principal components, explaining 65.02% and 12.75% of the variance, respectively. (B) Multidimensional Scaling (MDS) plot shows clear clustering of tumor and normal samples based on dissimilarity in the first two dimensions. (C) Density Plot highlights distribution differences, with peaks around a value of 24 for both groups. (D) Box Plot displays lower median protein expression levels in tumor samples compared to normal samples, indicating significant expression variation between the two groups.

Additionally, users can examine individual sample distributions using box plots and density plots, which highlight trends such as lower median protein expression in tumor samples compared to normal tissues (Figure 2C-D). Heatmaps with hierarchical clustering further summarize expression patterns, enabling researchers to identify distinct molecular signatures across cancer types (Supplementary Figure 1).

### 3.3 Differential Expressed Proteins and phosphoproteins

Users can conduct differential expression analysis to detect significantly altered proteins (DEPs) and phosphoproteins (DEPPs) across various cancer types or user-uploaded datasets. Using statistical methods such as limma, the platform calculates fold-change and adjusted p-values to pinpoint significant dysregulated features and further enables chooses to select different cutoffs. For example, analysis of CPTAC datasets revealed that FGFR2 and FGFR4 are downregulated in LUAD at protein and phosphoprotein levels, respectively. These results are consistent with previous findings of reduced FGFR4 expression in LUAD and its association with tumor stage and differentiation [50,51]. Additionally, FGFR4 interacts with EGFR, contributing to oncogenic signaling in LUAD, further highlighting its role in cancer progression [52]. Similarly, JAK2 is significantly downregulated at both protein and phosphoprotein levels, supporting its role in maintaining cancer stem cells and its contribution to radioresistance in colorectal cancer [53]. Meanwhile, carbonic anhydrase IX (CA9) is highly upregulated in CCRCC, reinforcing its utility as a diagnostic biomarker, although its occasional expression in other renal neoplasms limits its specificity [54]. In LSCC, TP53 is significantly upregulated, aligning with its frequent mutations in lung cancer and its established role in tumor progression and therapy resistance [55]. In contrast, EGFR is notably upregulated in ovarian cancer (logFC: -0.891, p-value: 5.07E-12), contributing to the ongoing uncertainty regarding its role as a prognostic biomarker [56].

In PDAC, CD44 is significantly downregulated (logFC: -1.19, p-value: 4.34E-14), despite its established role in tumor plasticity, invasion, and resistance to gemcitabine, suggesting potential therapeutic implications [57]. In UCEC, KRT17 is highly upregulated at both the proteome and phosphoproteome levels, reinforcing its involvement in tumor progression through the HIF-1α/VEGF signaling pathway, which enhances cancer cell migration and angiogenesis [58]. Conversely, WNT2, which has been implicated in UCEC progression, is actually down-regulated (logFC: -1.389, p-value: 7.46E-12), suggesting a more complex and context-dependent role in tumor biology [59]. PLK1 is strongly upregulated in LSCC, consistent with its known role in promoting tumor progression and cell cycle dysregulation [60]. In PDAC, transthyretin (TTR) shows significant upregulation (logFC: 1.148, p-value: 1.90E-12), aligning with its reported function in tumor metabolism and systemic response to pancreatic cancer [61]. Lastly, CHRNA3 is markedly upregulated in LUAD (logFC: 1.14, p-value: 1.26E-65), supporting its association with lung cancer susceptibility and its involvement in nicotine-related oncogenic pathways [62] (Table 2 and Supplementary Tables 1-2).

**Table 2.** Differentially Expressed Proteins and Phosphoproteins Across Cancer Types from CPTAC Data. This table presents differentially expressed proteins and phosphoproteins identified across multiple cancer types using CPTAC data. The table includes the log fold-change (logFC), average expression levels, and adjusted p-values for significant changes in expression.

| Type | Cancer | Gene | logFC | Average expression | Adjusted p-value |
| --- | --- | --- | --- | --- | --- |
| Proteome | LUAD | FGFR2 | -0.803 | 16.87 | 2.98E-62 |
|  | COAD | JAK2 | -1.154 | 20.851 | 6.94E-40 |
|  | CCRCC | CA9 | 1.922 | 25.478 | 1.70E-49 |
|  | LSCC | TP53 | 0.98 | 22.335 | 6.19E-13 |
|  | OV | EGFR | -0.891 | 23.651 | 5.07E-12 |
|  | PDAC | CD44 | -1.19 | 27.588 | 4.34E-14 |
|  | UCEC | KRT17 | 2.23 | 25.605 | 2.50E-10 |
|  | UCEC | WNT2 | -1.389 | 21.346 | 7.46E-12 |
| Phosphoproteome | COAD | JAK2 | -1.097 | 20.77 | 3.82E-41 |
|  | LSCC | PLK1 | 1.525 | 15.992 | 8.47E-77 |
|  | PDAC | TTR | 1.148 | 19.186 | 1.90E-12 |
|  | LUAD | CHRNA3 | 1.14 | 13.618 | 1.26E-65 |
|  | LUAD | FGFR2 | -1.769 | 13.77 | 4.86E-146 |
|  | LUAD | FGFR4 | -1.332 | 19.54 | 4.20E-77 |
|  | UCEC | KRT17 | 1.25 | 19.613 | 1.03E-14 |

Volcano plots with interactive features enable users to examine DEPs and DEPPs by displaying fold-change against statistical significance, facilitating easy recognition of crucial molecular components (Figure 3A). A boxplot illustrates variations in proteome or phosphoproteome expression levels for specific cancer data and can be further generated for one or multiple genes of interest. Heatmaps offer an additional perspective by grouping dysregulated features, exposing co-expression patterns that align with biological processes or pathways (Figure 3B).

**Figure 3.**
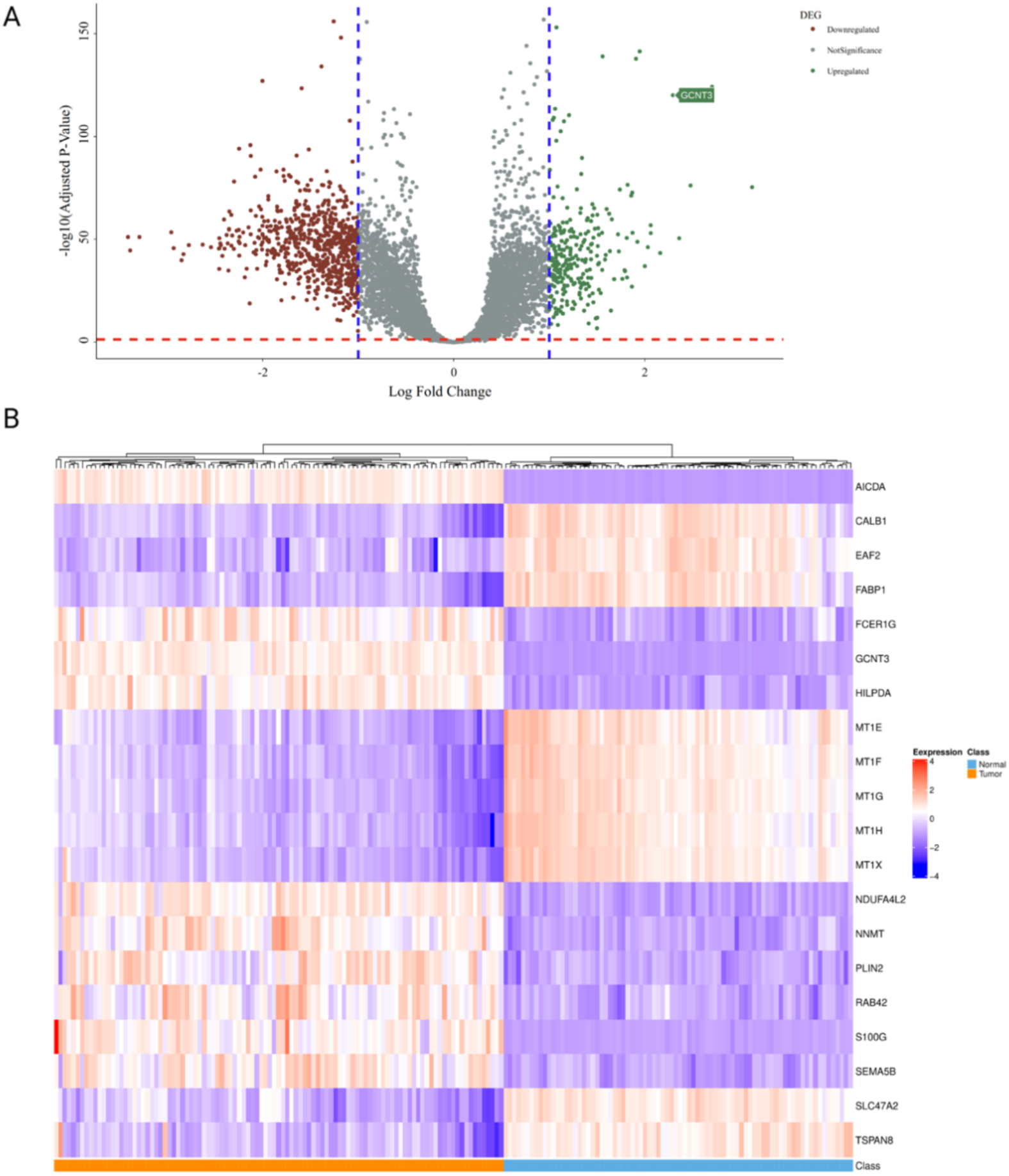
CCRCC differential proteomics visualization: (A) An interactive volcano plot displaying -log10(adjusted P-value) on the y-axis and log2 fold change on the x-axis, highlighting differential expression patterns. Vertical dashed lines represent log fold change thresholds, while the horizontal dashed red line denotes the significance threshold. Hovering over data points reveals gene symbols, enabling detailed examination of significant differentially expressed proteins (DEPs) and phosphoproteins (DEPPs). (B) A heatmap of top user-selected genes, with genes represented as rows and samples as columns. Hierarchical clustering at the top groups samples, uncovering co-expression patterns aligned with biological processes or pathways.

### 3.4 Enrichment analysis of DEPS and DEPPS, pathway visualization, PPI and Drug interaction analysis

OncoProExp integrates comprehensive gene enrichment analysis implemented using the gProfileR package for differentially expressed proteins (DEPs) and phosphoproteins (DEPPs). This allows users to explore biological pathways and functional annotations across multiple databases, including GO, KEGG, REACTOME, TF, miRNA, HPA, CORUM, and WikiPathways. Users can perform enrichment analysis separately for upregulated, downregulated, or both DEPs/DEPPs, thus tailoring the analysis to specific research questions. A unique aspect of OncoProExp is its KEGG pathway enrichment analysis, which maps dysregulated proteins onto cancer-relevant pathways. Using the pathview tool, users can visualize up- and downregulated genes directly within KEGG pathway graphs, providing an intuitive and interactive way to interpret pathway-level changes in the context of cancer biology. For example, in PDAC datasets, OncoProExp highlighted the activation of the pancreatic secretion pathway, reflecting its role in tumor development (Figure 4A). Through the Pathview package, enriched pathways can be visualized with up- or down-regulated proteins clearly mapped onto KEGG pathway graphs.In PDAC, OncoProExp identified the upregulation of ITPR3 (Figure 4B), a key regulator of intracellular Ca²⁺ release, which enhances calcium signaling to promote tumor cell proliferation, survival, and endoplasmic reticulum stress [63].

**Figure 4.**
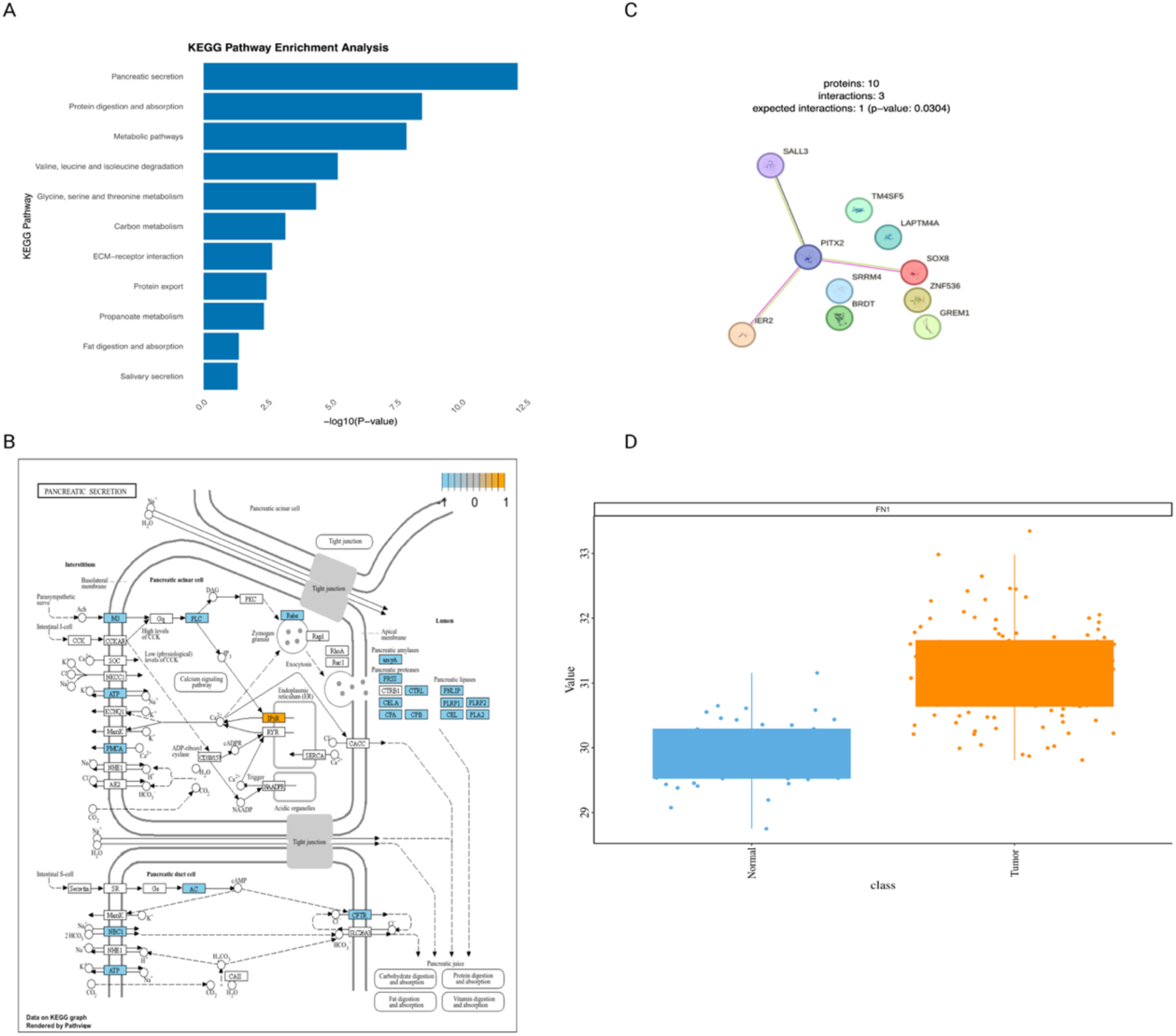
Analysis of Pathways, Protein Interactions, and Drug Targets in Pancreatic Ductal Adenocarcinoma (PDAC) using OncoproExp. (A) KEGG pathway analysis identifies pancreatic secretion as the most enriched pathway in PDAC datasets. (B) Pathview visualization shows ITPR3 upregulation, indicating increased calcium signaling, while PMCA and ATP-related genes are downregulated. (C) The Protein-Protein Interaction (PPI) network, generated using OncoProExp in the LUAD proteome dataset, reveals PITX2 as a central gene in a four-connection network (p-value = 0.0304). (D) Boxplot analysis demonstrates significant FN1 overexpression in pancreatic ductal adenocarcinoma (PDAC) compared to normal tissues. Data from CancerDrugs_DB shows that Dacarbazine (DrugBank ID: DB00851) targets FN1, which is notably upregulated in PDAC.

Meanwhile, PMCA and ATP-related gene downregulation indicated a reduced calcium-efflux capacity and metabolic strain, reflecting dysregulated calcium homeostasis. Consistent with these observations, Richardson et al. demonstrate that PDAC cells rely on glycolytic ATP to power PMCA, and disrupting this localized energy source selectively impairs calcium pumps—causing cytotoxic Ca²⁺ overload without depleting global ATP [64]. These findings demonstrate the tool’s ability to uncover critical molecular mechanisms and provide pathway-level insights into cancer biology.

Users can perform protein-protein interaction (PPI) analysis using OncoProExp to identify key interaction hubs and explore the connectivity among dysregulated proteins. For example, analysis of the protein-protein interaction (PPI) network for the top 10 differentially expressed proteins (DEPs) in the LUAD proteome, using CPTAC data, resulted in four interactions, higher than the expected interaction of one (p-value: 0.0304) (Figure 4C). The resulting network highlighted PITX2, SALL3, SOX8, and IER2 as crucial nodes. PITX2 functions as an oncogene in LUAD by transcriptionally upregulating WNT3A, thereby activating the Wnt/β-catenin signaling pathway to promote tumor progression [65]. In contrast, SALL3 serves as a tumor suppressor, with its epigenetic silencing through promoter hypermethylation is linked to poor prognosis, aggressive tumor behavior, and reduced disease-free survival in HNSCC [66]. Similarly, SOX8 contributes to cancer stem-like cell maintenance and tumor progression in triple-negative breast cancer [67]. The immediate early gene IER2 further underscores its significance in cancer biology by enhancing tumor cell motility and invasiveness, facilitating metastasis, and correlating with poor survival in colorectal cancer patients, making it a promising prognostic biomarker and therapeutic target [68]. Thus, the interconnections observed among these proteins point to a functional unit that may contribute to vital cancer-related processes, including cellular differentiation, tumor growth, and metastasis.

Users can further explore drug target information integrated from CancerDrugs_DB, linking dysregulated proteins to potential therapeutic interventions. For instance, a boxplot analysis of FN1 in PDAC revealed its significant upregulation in tumor samples compared to normal tissues. FN1, identified as a target of the chemotherapeutic agent Dacarbazine, highlights the translational potential of OncoProExp in connecting molecular alterations to actionable drug targets (Figure 4D). These findings underscore the utility of OncoProExp in connecting molecular changes to biological functions, supporting hypothesis generation, and targeted experimental design.

### 3.5 Pan-cancer analysis

The pan-cancer module of OncoProExp allows users to investigate differentially expressed proteins (DEPs) and phosphoproteins (DEPPs) across various cancer types using the CPTAC datasets. This feature enables the comparison of expression levels between normal and tumor samples for multiple cancers with the ability to generate box plots for visual representation. For instance, users can examine EPCAM expression in lung adenocarcinoma (LUAD), ovarian cancer (OV), and clear cell renal cell carcinoma (CCRCC), revealing distinct patterns between tumors and normal tissues (Supplementary Figure 2). In LUAD, EPCAM is frequently overexpressed, though its prognostic relevance remains uncertain [69,70], whereas in CCRCC, EPCAM often correlates with favorable tumor features [71]. In contrast, ovarian cancer overexpresses EPCAM in chemoresistant cancer stem cell–like populations, contributing to poorer clinical outcomes [72]. The platform also provides a comprehensive table of DEPs and DEPPs for all CPTAC cancer types, filtered for significance (FDR < 0.05 for DEPs/DEPPs and *p* < 0.05 for survival biomarkers). This consolidated view offers insights into dysregulated proteins and phosphoproteins, helping identify shared and cancer-specific molecular signatures. Furthermore, the pan-cancer module facilitates the exploration of survival biomarkers, allowing researchers to assess the prognostic significance of specific proteins across multiple cancer types.

### 3.6 Feature Selection

The selected features, derived through a two-step process combining Random Forest importance scores and MAD-based filtering, demonstrated a robust discriminatory power. The t-SNE projections revealed distinct clustering of tumor and normal samples across all cancer types in both proteome (Figure 5A) and phosphoproteome (Figure 5B) datasets. Clear separation between tumor and normal groups, as well as distinct cancer-specific clusters, highlights the biological relevance of the top 250 proteins (Supplementary file 2) and 100 phosphoproteins (Supplementary file 3). These findings confirm that the selected features effectively capture meaningful variations for accurate classification in downstream machine learning models.

**Figure 5.**
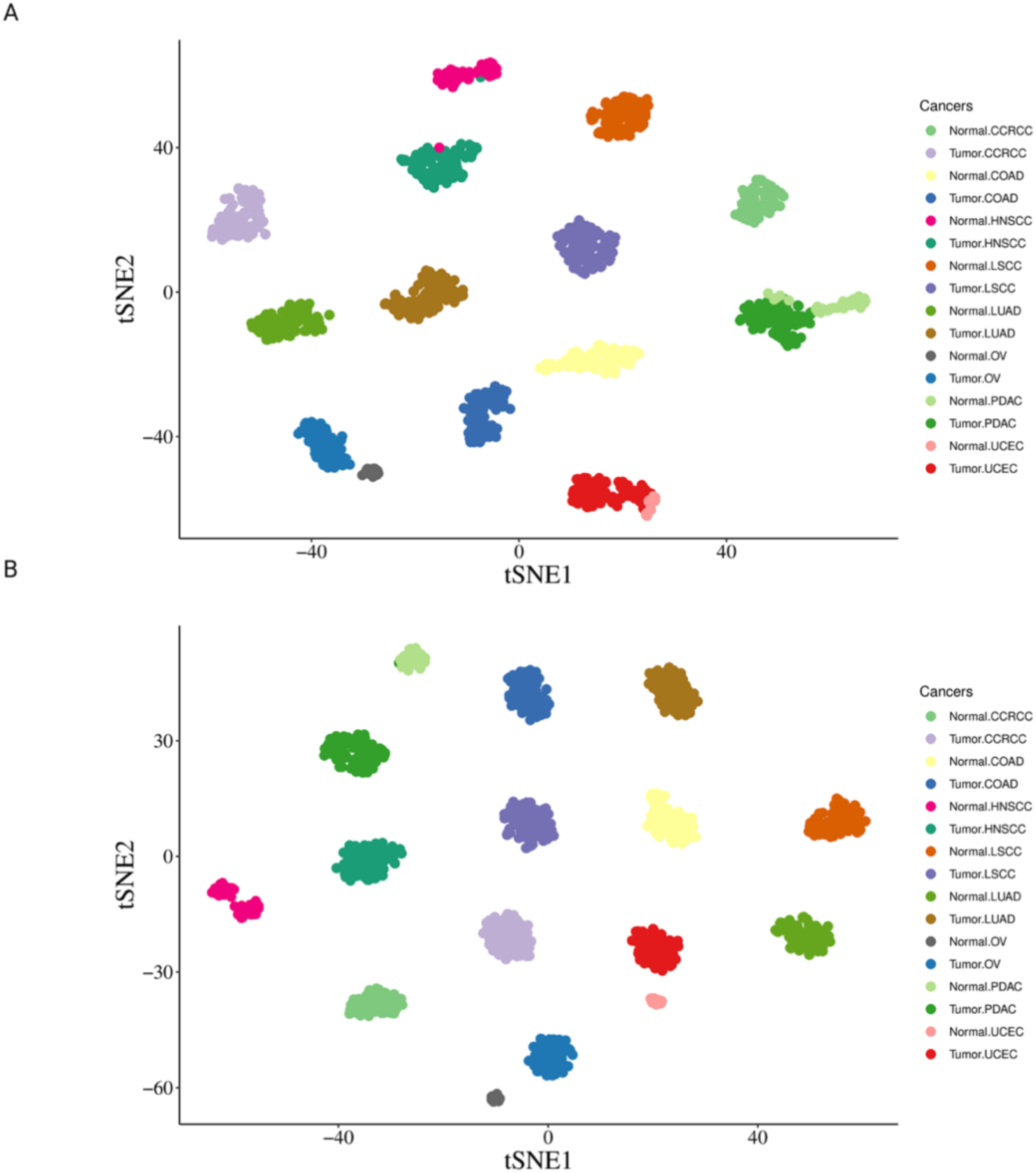
t-SNE analysis of (A) proteome and (B) phosphoproteome data, illustrating separation between tumor and normal samples across multiple cancer types (CCRCC, COAD, HNSCC, LUAD, LSCC, OV, PDAC, and UCEC). Each point is colored by cancer type and sample origin, revealing distinct clusters for different cancers and a clear separation between tumor and normal samples within each cancer. The clustering patterns indicate significant differences in protein and phosphoprotein expression profiles, underscoring the potential of both data types to effectively distinguish tumor from normal tissues.

### 3.7 AI-Based Predictive Models and Their Performance

OncoProExp integrates advanced machine learning algorithms, including Support Vector Machines (SVM), Random Forests (RF), and Artificial Neural Networks (ANN), to predict cancer types from proteomic and phosphoproteomic data. Users can train these models in both multi-cancer and single-cancer modes, thereby achieving high accuracy and robustness across the test and validation sets.

#### 3.7.1 Multi-Cancer Mode

In the multi-cancer mode for proteomics data, the SVM model achieved perfect classification on the test set, with all metrics (accuracy, precision, sensitivity, F1 score, specificity, and AUC) reaching 1.0, reflecting flawless performance across all cancer types (Supplementary Tables 3-4). On the validation set, the SVM model also performed exceptionally, demonstrating high accuracy and excellent scores in precision, sensitivity, F1 score, and specificity, particularly for HNSCC and PDAC, and maintaining an AUC of 0.996 for all classes (Supplementary Tables 5-6).

The RF model showed similarly high performance on the test set, achieving perfect classification for all cancer types with an accuracy, precision, sensitivity, F1 score, and specificity of 1.0 for most classes (Supplementary Tables 7-8). Minor variations were observed in Normal PDAC and Tumor PDAC with slight reductions in accuracy and F1 score, but still demonstrating near-perfect results. On the validation set, the RF model maintained high performance, with accuracy, precision, sensitivity, F1 score, and AUC values reflecting robustness in classifying both HNSCC and PDAC samples (Supplementary Tables 9-10).

The ANN model also exhibited excellent performance across the test set, achieving perfect scores in all metrics for most cancer types and maintaining high precision and recall (Supplementary Tables 11-12). On the validation set, the ANN model’s results mirrored those of the RF model, with high accuracy and robust performance metrics for both HNSCC and PDAC, again demonstrating high consistency and reliability (Supplementary Tables 13-14).

#### 3.7.2 Single-Cancer Mode

In single-cancer classification mode, the SVM, RF, and ANN models demonstrated highly accurate performance across HNSCC and PDAC. For HNSCC, all three models (SVM, RF, and ANN) achieved identical performances, with 99.4% accuracy, 100% precision, and 98.4% sensitivity, resulting in an F1 score of 0.992. The AUC for all models was 0.995, highlighting their strong discriminatory power (Supplementary Tables 15-16). For PDAC, the SVM, RF, and ANN models performed equally well, each achieving an accuracy of 97.2%, with 96% precision, 96% sensitivity, and an F1 score of 0.96. The AUC for all models was 0.969, further emphasizing their robust classification ability (Supplementary Tables 17-18). Overall, all three models exhibited consistent and high performance in distinguishing tumor and normal samples in both HNSCC and PDAC, with HNSCC classification reaching near-perfect accuracy and PDAC classification maintaining strong predictive reliability.

#### 3.7.3 Phosphoproteome Data

The evaluation of machine learning models on phosphoproteome data demonstrated exceptional classification performance in both multi-cancer and single-cancer classification modes. In multi-cancer classification mode, which included CCRCC, COAD, HNSCC, LSCC, LUAD, OV, PDAC, and UCEC, the SVM, RF, and ANN models achieved perfect classification accuracy across all cancer types. The confusion matrices for these models showed no misclassifications, with each model accurately distinguishing between normal and tumor samples. The key performance metrics, including accuracy, precision, sensitivity, F1 score, specificity, and AUC, all reached 1.0, underscoring the reliability of the models (Supplementary Tables 19-24).

Similarly, in single-cancer classification mode for CCRCC, all three models (SVM, RF, and ANN) again achieved perfect differentiation between normal and tumor samples, as confirmed by the confusion matrices (Supplementary Table 25). The performance metrics were uniformly optimal (accuracy = 1.0 across all models), reinforcing the models’ ability to classify CCRCC with complete accuracy (Supplementary Table 26). Overall, the analysis confirms that the models excel in classifying cancer and normal samples using proteome and phosphoproteome data, showcasing their robustness and precision in both multi-cancer and single-cancer settings.

### 3.8 Model Explanation Using SHAP

To enhance the interpretability of AI-based predictions, OncoProExp incorporates SHapley Additive exPlanations (SHAP). Users can explore the contributions of individual proteins and phosphoproteins to model predictions, gaining insights into the molecular features driving cancer classification.

#### 3.8.1 Multi-Cancer Model

In the multi-cancer model for proteome data(Figure 6A), genes such as ITGA7 and LILRB5 showed mixed impacts, with ITGA7 having a broader influence and MYO1C significantly boosting the model. In the phosphoproteome data (Figure 6B), XPO4 and COLGALT1 displayed varied effects, with XPO4 having a wider influence, and SYN1 showing a strong positive impact. Other genes, such as SEPSECS, MCCC2, and PLIN4, contributed both positively and negatively, while GYPC and LRP1 exhibited diverse impacts, with LRP1 leaning positively. FAM83F and CD34 showed mixed effects, with CD34 exhibiting a broader range.

**Figure 6.**
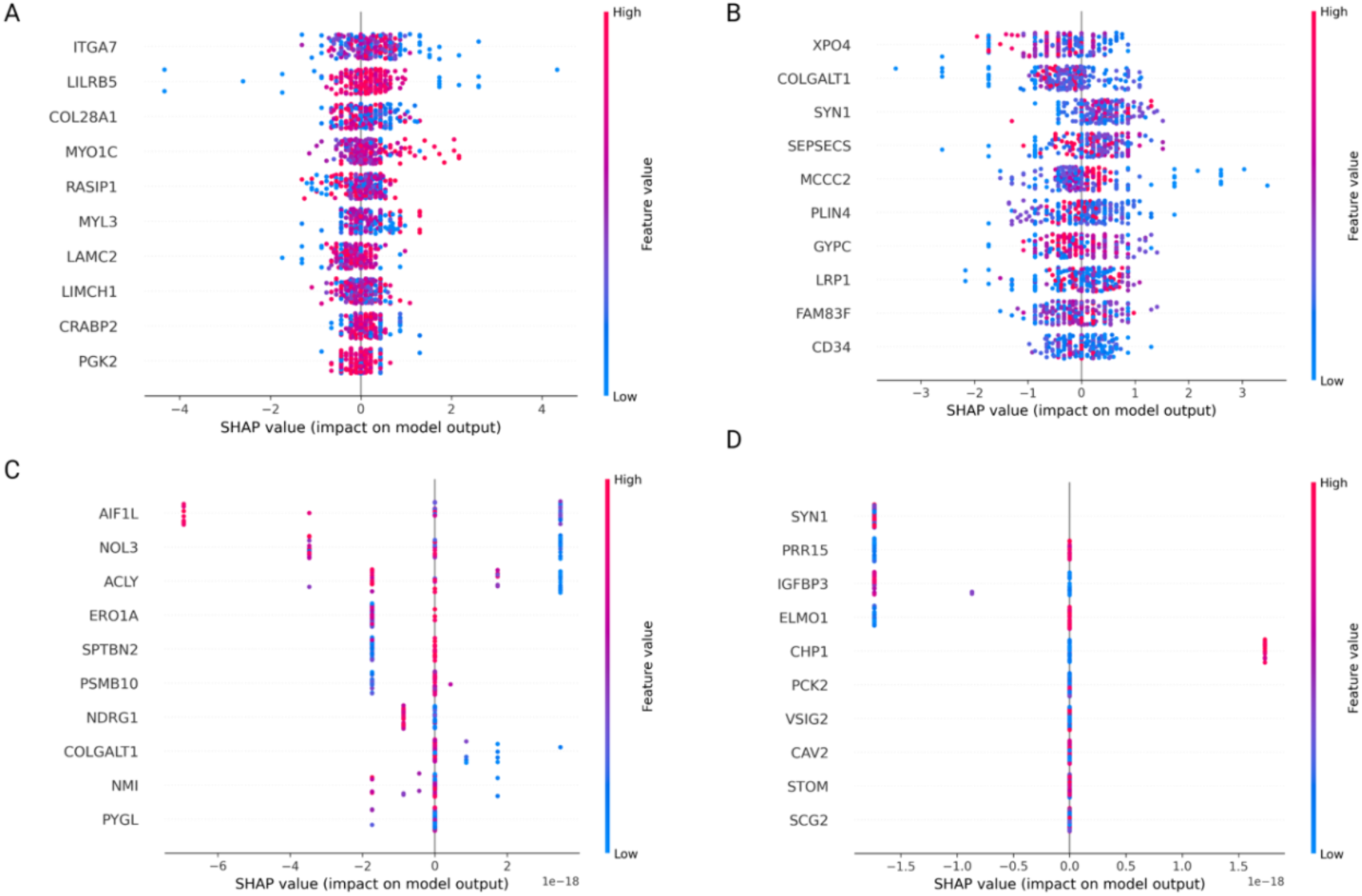
Interpretation of model predictions using SHAP values. SHAP summary plots show the impact of top proteins (A) and phosphoproteins (B) on model predictions in multi-cancer mode. Each point represents a sample, with its x-position indicating the SHAP value for a given feature (impact on model output), while the vertical axis lists the features. Its color denotes the feature value (red: high, blue: low). The spread of points along the x-axis indicates the range of a feature’s impact. These plots show that most features contribute to both increases and decreases in the model’s predicted values. Panels (C) and (D) highlight features with more consistent negative or positive impacts in single-cancer mode, respectively.

#### 3.8.2 Single-Cancer Model

In the single-cancer model for CCRCC proteome data (Figure 6C), AIF1L and NOL3 showed strong negative impacts on the model output. A similar negative trend, though less pronounced, was observed for ACLY, ERO1A, and NDRG1. For the same cancer type in phosphoproteome data (Figure 6D), genes like SYN1 and IGFBP3 showed strong negative impacts, while CHP1 exhibited a positive impact. Higher expression levels (represented by pink for proteome and red for phosphoproteome) correlated with stronger impacts, highlighting the diverse roles of genes in model performance.

### 3.9 Survival analysis

OncoProExp allows users to identify biomarkers with prognostic value across various cancer types through its extensive survival modeling capabilities. For instance, users can investigate the effects of CA12 proteome expression in UCEC or MGMT proteome expression in GBM by creating Kaplan-Meier plots and determining hazard ratios (HR). The system automatically selects the optimal expression threshold for each biomarker, ensuring balanced group comparisons and accurate HR calculations. In the case of CA12 in UCEC, elevated expression was linked to significantly poorer survival outcomes (HR: 0.422, p-value = 0.01), aligning with its established role in facilitating tumor progression via pH regulation and hypoxia adaptation [73]. Likewise, reduced MGMT expression in GBM was associated with decreased survival (HR: 1.8, p-value =0.03), reflecting its involvement in treatment resistance [74,75]. These results, depicted through user-friendly Kaplan-Meier graphs (Supplementary Figure 3), underscore the clinical significance of these biomarkers and their potential as therapeutic targets. By allowing users to control for variables such as age or tumor stage, OncoProExp enables the discovery of actionable insights and customization of survival analyses to address specific research questions.

### 3.10 Comparison of OncoproExp with similar available tools

OncoProExp sets itself apart from other cancer proteomics analysis tools by integrating machine learning (ML) with statistical approaches. Unlike PhosMap [76], cProSite [77], TCPA [78], CPPA [79], UALCAN [80], and iProPhos [81] which primarily use basic statistical methods, OncoProExp employs advanced ML classifiers such as Random Forest (RF), Support Vector Machines (SVM), and Artificial Neural Networks (ANN), in combination with limma for differential expression analysis. A unique feature of OncoProExp is its incorporation of SHAP (SHapley Additive exPlanations) for CPTAC data, providing explainable AI insights which is absent in other tools. Moreover, OncoproExp excels in comprehensive dimensionality reduction (PCA, MDS, UMAP), pathway enrichment (GSEA, KEGG, Reactome, PPI networks), and advanced missing-value imputation techniques, surpassing existing tools. OncoProExp also offers robust survival analysis, including univariate and multivariate Cox models and Kaplan-Meier plots, and evaluates therapeutic relevance by cross-referencing dysregulated proteins with drug databases like DrugBank and the Cancer Drugs Database. While UALCAN and iProPhos provide cancer database integration and user data upload, OncoProExp enhances accessibility and reproducibility through its web-based interactive dashboard and Docker functionality. In essence, OncoProExp emerges as a versatile and powerful tool for cancer proteomics, blending ML-driven analytics with user-friendly features, making it the preferred choice for researchers seeking advanced, interpretable, and comprehensive cancer data analysis. Table 3 presents a comprehensive comparison of OncoproExp’s capabilities with those of current Phosphoproteome/proteome tools.

**Table 3.** Comprehensive comparison of OncoproExp’s with those of current Phosphoproteome/proteome tools.

| Feature | OncoProExp | PhosMap | TCPA | cProSite | UALCAN | iProPhos |
| --- | --- | --- | --- | --- | --- | --- |
| Machine Learning vs. Basic Statistical Methods | ML + Stats (advanced classifiers (RF, SVM, ANN) plus limma) | Stats only (limma, ANOVA, etc.) | Stats only (correlation, t-test, univariate K-M) | Stats only (basic group comparisons, correlations) | Stats only (basic group comparisons, correlation, survival analyses) | Stats only (differential expression, correlation, KSEA) |
| SHAP / Explainable AI Integration | Yes (Single and pan-cancer) | No | No | No | No | No |
| Dimensionality Reduction | Yes (PCA, MDS, UMAP) | Yes (PCA, t-SNE, UMAP) | No (clustered heatmap, correlation network) | No (heatmaps, correlation) | No (primarily boxplots, correlation scatter, survival curves) | No (focus on differential expression and correlation analyses) |
| Pathway / Network Enrichment | Yes (GSEA, KEGG, Reactome, PPI network) | Limited (KSEA, motif-x; no direct KEGG/Pathview) | No (only RPPA-based correlations) | No | No | Yes (Over-Representation, GSEA, and PPI network analyses) |
| Advanced Missing-Value Imputation | Yes (Using percentage of NAs, RF, mean, medain, mod) | Yes (Min, BPCA, LLS, kNN, etc.) | No | No | No | Yes (filters >50% missing then applies kNN) |
| Survival Analysis | Yes (univariate and multivariable Cox, Kaplan-Meier plot) | Yes (multivariable CoxPH available) | No (univariate Kaplan-Meier only) | No | Yes (univariate Kaplan-Meier with some multivariable options) | Yes (Kaplan-Meier/log-rank tests with user-specified cutoffs) |
| Plotting & Visualization | Volcano plots, Heatmaps, PCA/UMAP/MDS, Scree plot, Box and density plots, GSEA plot, Enrichment plot, KEGG image, PPI network, Boxplots for drug-targetable genes, Kaplan-Meier curves, SHAP | Volcano plots, Heatmaps, Correlation scatter, PCA/t-SNE/UMAP, Survival curves | Clustered heatmap, Correlation network, Basic survival plots | Boxplots, Barplots, Heatmaps, Tumor vs. Normal fold-change plots | Boxplots, Correlation scatter, Kaplan-Meier curves, Pan-cancer comparisons | Volcano plots, Boxplots, Correlation scatter, KSEA barplots, Kaplan-Meier curves, PPI network maps |
| Therapeutic Relevance Evaluation | Yes (dysregulated proteins) | No | No | No | No | Partial (identifies potential drug targets and upstream kinases, but |
|  | are cross-referenced with DrugBank and the Cancer Drugs Database; box plots illustrate differential expression of drug-targetable genes) |  |  |  |  | does not explicitly cross-reference with public drug databases) |
| User Data Upload | Yes (Allows upload of proteomics/phosphoproteomics data along with their metadata) | No | No | No | No | Yes (Allows upload of custom proteomics/phosphoproteomics data) |
| Cancer Database Integration | Yes (integrates CPTAC proteomics/phoproteomics) | No (requires user-provided data) | Yes (built on TCGA RPPA data plus CCLE cell lines) | Yes (CPTAC proteomics/phosphoproteomics and GDC/TCGA mRNA) | Yes (integrates TCGA transcriptome, miRNA, lncRNA, promoter methylation and CPTAC proteomics) | Yes (integrates CPTAC proteome & phosphoproteome from 12 cancer types plus matched transcriptome) |
| User Interface (Web vs. Local vs. CLI) | Web + Docker (interactive dashboard) | Local (Docker container, R-based) | Web-based portal | Web-based portal | Web-based portal | Web-based (R Shiny), no Docker |
| Docker Functionality | Yes | Yes | No | No | No | No (R Shiny web server; no Docker image) |
| License / Source Code | Open source (Entire code on GitLab link) | Open source (GitHub: BADD-XMU/Phos-Map) | Free access (hosted; not fully open-sourced) | Free (hosted by NCI-CCR; no open-source statement) | Free (hosted by UAB; partial code available on GitHub) | Free (hosted by Zhejiang University; code to be available on GitHub upon publication) |

## Discussion

This study presents OncoProExp, a web-based unified platform designed to facilitate analysis of proteomic and phosphoproteomic data in cancer research. The platform integrating mass spectrometry (MS)-based datasets with advanced computational tools, OncoProExp enables researchers to explore dysregulated proteome and phosphoproteomes and their associations with cancer progression and therapeutic targets. This platform’s user-friendly interface eliminates the need for advanced bioinformatics expertise, making high-throughput proteomics accessible to a broad scientific audience. OncoProExp supports multilayered analyses by integrating clinical, proteomic, and phosphoproteomic data, facilitating biomarker discovery and translational research. It combines robust preprocessing workflows, statistical analyses, and ML models, offering key functionalities, such as differential expression analysis, pathway enrichment, and AI-driven predictive modeling. To address critical gaps in existing tools, OncoProExp serves as a comprehensive resource for cancer biomarker discovery, therapeutic target identification, and precision oncology.

Cancer biomarkers have been extensively studied using genetic and epigenetic approaches including circulating miRNAs and DNA methylation [82–87]. However, large-scale proteomic studies remain underutilized for pan-cancer analysis. Notable efforts include the work of Álvez et al., who employed machine learning for pan-cancer blood proteome profiling using proximity extension assays. However, large-scale proteomic studies remain underutilized for pan-cancer analysis. Notable efforts, such as those by Álvez et al., who utilized machine learning techniques to analyze pan-cancer blood proteomes using proximity extension assays [3], and Gonçalves et al., who mapped proteomic landscapes across 949 human cancer cell lines [88], highlighted the potential of proteomics in cancer research. Despite these advances, existing tools such as the like ProteoCancer Analysis Suite (PCAS) [89], cProSite, TCPA, CPPA Click or tap here to enter text., UALCANClick or tap here to enter text., LinkedOmics [90], cBio Cancer Genomics Portal [91], CancerProteome [11], BioLadder [92], OSppc [93], PhosMap, phosphoproteome prediction[94] and iProPhos have primarily focused on data visualization and basic analysis. Many existing tools lack robust data-processing pipelines, user-provided data integration, customizable visualizations, and AI-driven predictive modeling capabilities. They frequently omit crucial elements, such as functional analysis, drug target evaluation, and pathway-focused visualizations, which are essential for extracting clinical insights and advancing precision oncology. Moreover, platforms such as iProPhos prioritize PTM prediction, but fail to incorporate clinical correlations. OncoProExp addresses this gap by linking phosphosites (e.g., PLK1 in LSCC) to patient survival and therapeutic targets through cross-database interoperability, including DrugBank integration, thereby enhancing its clinical relevance.

OncoProExp distinguishes itself by integrating pan-cancer proteomic analyses with advanced ML algorithms such as SVMs, RFs, and ANNs. The platform facilitates predictive modeling, biomarker discovery, and pathway analysis, whereas SHAP improves model interpretability, providing insights into the molecular mechanisms driving cancer. These features allow researchers to move beyond traditional profiling and leverage AI for predictive modeling, therapeutic target identification with greater precision and efficiency, and pathway analysis on a single platform. OncoProExp outperforms existing methods in classification accuracy (up to 99.4% in HNSCC and PDAC) and provides SHAP-based model interpretability—features absent in many existing tools. Additionally, OncoProExp supports functional annotation and drug repurposing through enrichment analysis and protein-protein interaction networks. Linking dysregulated proteins to curated drug databases enables the identification of potential therapeutic targets and drug candidates [39,40]. This is particularly valuable for precision oncology, where targeted therapies rely on the identification of actionable molecular alterations. Furthermore, OncoProExp allows pan-cancer analysis, facilitating the identification of shared oncogenic drivers and tumor-specific proteomic signatures.

A key strength of OncoProExp is its ability to perform differential expression analysis, enabling the identification of cancer-specific proteins and phosphoproteins. This capability, combined with functional enrichment analyses such as KEGG pathway mapping and GSEA, provides users with deeper insight into cancer progression. Moreover, the platform’s interactive visualization tools, such as PCA, hierarchical clustering heatmaps, and UMAP, further enhance data exploration, interpretation, and hypothesis generation. The inclusion of CPTAC datasets enhances cross-cancer comparisons, enabling the identification of shared and unique molecular signatures that are critical for therapeutic targeting and patient stratification. OncoProExp incorporates phosphoproteomic data to offer a unique perspective on PTMs, which often play key roles in oncogenesis and treatment resistance. Furthermore, OncoProExp supports translational applications by integrating with cancer drug databases [39,40] and survival analysis tools, enabling researchers to connect dysregulated proteins to established therapies and assess their clinical relevance. The predictive modeling capabilities of the platforms extend to both pan-cancer and single-cancer levels, supporting the full pipeline from biomarker discovery to clinical application. OncoProExp’s user-friendly Shiny-based interface and Docker-based deployment ensure accessibility to researchers with limited bioinformatics expertise, broadening its applicability in both basic and clinical research settings. OncoProExp promotes collaboration through its data-sharing features, allowing researchers to export and share analysis results easily. The platform’s standardized data processing pipeline ensures reproducibility across different research groups, facilitating multi-institutional studies, and accelerating the pace of cancer proteomics research.

## Limitations

Despite its strengths, the OncoProExp has several limitations. Its reliance on CPTAC datasets restricts the inclusion of diverse data types, thus limiting the inclusion of other PTMs, such as glycosylation and acetylation. In addition, large dataset processing may slow the performance of network-based analyses or ML-driven predictions. Furthermore, the platform facilitates hypothesis generation, experimental validation remains essential to confirm the clinical utility of the identified biomarkers and therapeutic targets.

To address these limitations, future updates will expand OncoProExp’s analytical scope beyond CPTAC data by integrating additional multiomics datasets for broader comparative studies. Furthermore, the platform will incorporate advanced data mining techniques to extract experimental validation of proteins from the literature. Using text-mining algorithms and natural language processing (NLP), OncoProExp will systematically retrieve, and curate validated protein interactions, functional annotations, and biomarker evidence from peer-reviewed studies, thereby enhancing the reliability of protein discoveries and their clinical relevance. Additionally, we are optimizing the platform’s performance for large-scale datasets by implementing advanced data compression and distributed computing techniques, improving efficiency and scalability while maintaining robust analytical capabilities.

## Conclusion

OncoProExp fills a critical gap in proteomics research by integrating computational methods, AI-driven predictive modeling, and interactive visualizations into a single platform. Its application to CPTAC datasets demonstrated its potential for uncovering novel insights into cancer biology, paving the way for improved diagnostics and targeted therapies. By incorporating user feedback and evolving technological advancements, OncoProExp is expected to remain a valuable tool in cancer proteomics research, enabling researchers to translate complex datasets into actionable insights for precision oncology research.

## Data availability

Data for developing the tool are downloaded from CPTAC consortium and can be accessed from the following link: https://proteomics.cancer.gov/programs/cptac

The code is available from the following **link** https://gitlab.cs.ut.ee/edris/oncoproexp.git

## Author contributions

Edris Sharif Rahmani: Developed software, performed data curation and validation, and drafted the original manuscript. Prakash Lingasamy: Contributed to the original draft, conducted data interpretation, and participated in manuscript review and editing. Soheyla Khojand: Assisted in data interpretation and contributed to manuscript review and editing.

Ankita Lawarde: Performed formal analysis and contributed to manuscript review and editing. Sergio Vela Moreno: Assisted in manuscript review and editing. Andres Salumets: Provided funding acquisition and contributed to manuscript review and editing. Vijayachitra Modhukur: Led conceptualization and supervision, provided resources, conducted formal analysis, and contributed to the original draft.

## Acknowledgements

We would like to thank the HPC team from the University of Tartu for deploying the application, maintaining the system, and providing technical support. We also appreciate the CPTAC consortium for making their data publicly available. The graphical abstract was created using BioRender.

## Funding

The research was supported by the Horizon Europe grant (NESTOR, no. 101120075) and Estonian Research Council (grant no PRG1076).

## Conflict of interest statement

None declared.

## Notes

### Competing Interest Statement

The authors have declared no competing interest.

