## Supplementary file for "OncoProExp: An Interactive Shiny Web Application for Comprehensive Cancer Proteomics and Phosphoproteomics Analysis"

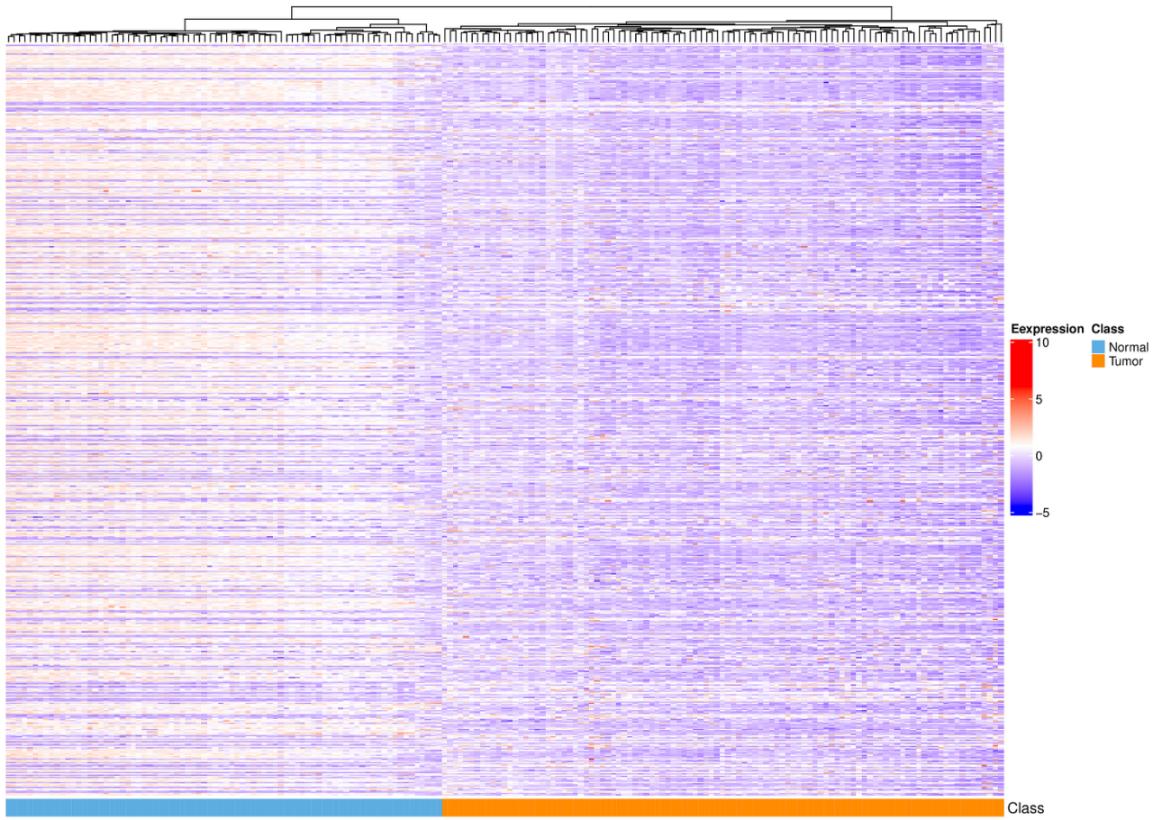
**Supplementary Figure 1: Heatmap of the CCRCC Proteome Dataset with Hierarchical Clustering** This heatmap visualizes the expression patterns of proteins in the CCRCC dataset, with hierarchical clustering applied to group samples. Tumor and normal samples are well-separated into distinct clusters, showcasing unique molecular signatures between the two groups. Heatmaps with hierarchical clustering provide a clear summary of expression patterns, enabling researchers to identify distinct molecular signatures across cancer types.

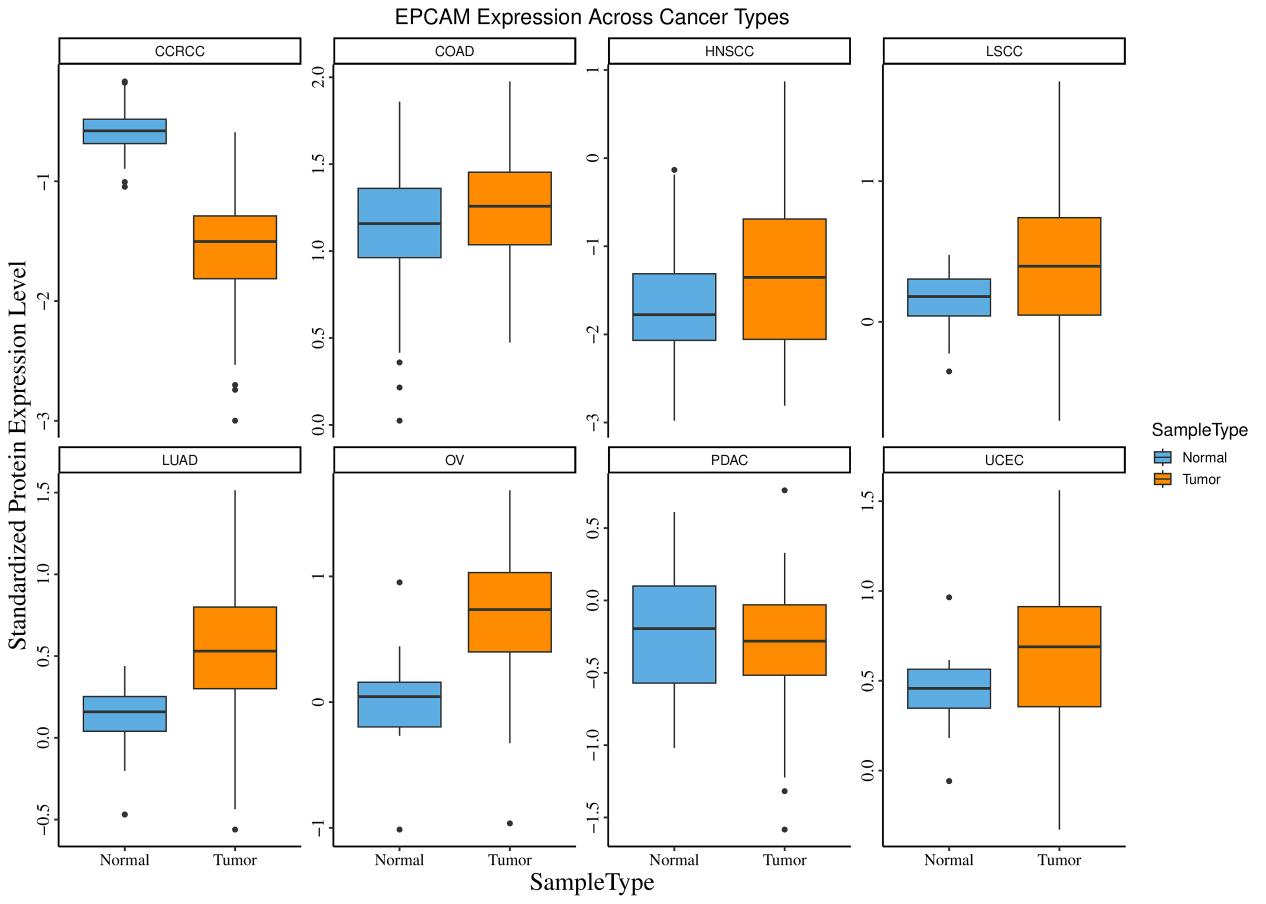

Supplementary Figure 2: EPCAM Expression Across Cancer Types
This figure highlights standardized protein expression levels of EPCAM in various cancer types, including LUAD, OV, and CCRCC, comparing tumor and normal tissues. Distinct expression patterns between tumor and normal samples are evident, providing insights into EPCAM's differential regulation across these cancer types.

### **Supplementary Table 1: Differentially Expressed Proteins Across Cancer Types from CPTAC Data.** This table presents the significantly differentially expressed proteins in various cancer types, including CCRCC, HNSCC, LSCC, PDAC, and UCEC, based on CPTAC proteomic data. The log fold-change (logFC), average expression levels, and adjusted p-values indicate the magnitude and statistical significance of expression changes.

| Cancer | Gene | logFC | Average expression | Adjusted p-value |
| --- | --- | --- | --- | --- |
| CCRCC | VEGFA | 1.25 | 23.003 | 1.58e-33 |
| CCRCC | CXCR4 | 0.911 | 17.959 | 2.5e-43 |
| HNSCC | NFE2L2 | 0.935 | 15.098 | 4.92e-47 |
| LSCC | KIT | -1.072 | 24.602 | 6.69e-29 |
| PDAC | ALB | -0.925 | 21.86 | 0.9e-5 |
| PDAC | EGFR | -0.891 | 23.651 | 5.07e-12 |
| PDAC | TTR | -0.863 | 28.274 | 0.2e-5 |
| UCEC | PTEN | -0.871 | 23.324 | 1.69e-7 |
| UCEC | ENPP3 | -0.843 | 23.015 | 3e-2 |

### **Supplementary Table 2: Differentially Expressed Phosphoproteins Across Cancer Types from CPTAC Data.** This table summarizes the significantly differentially expressed phosphoproteins across multiple cancer types, highlighting key regulatory proteins involved in oncogenic pathways. The table includes log fold-change (logFC), average expression levels, and adjusted p-values to represent statistically significant changes in phosphorylation levels.

| Cancer | Gene | logFC | Average expression | Adjusted p-value |
| --- | --- | --- | --- | --- |
| CCRCC | CA9 | 1.325 | 16.824 | 2.28e-74 |
| CCRCC | CD274 | 0.928 | 18.661 | 6.15e-32 |
| CCRCC | IRF5 | 0.903 | 17.647 | 6.43e-43 |
| HNSCC | DNMT3B | 0.903 | 14.32 | 1.35e-108 |
| LSCC | TP53 | 1.375 | 18.074 | 1.8e-47 |
| LSCC | FGFR3 | 0.847 | 16.096 | 2.19e-44 |
| LSCC | FGFR4 | -1.633 | 19.668 | 8.98e-75 |
| LSCC | ERBB4 | -0.812 | 15.702 | 2.59e-36 |
| PDAC | TTR | 1.148 | 19.186 | 1.9e-12 |
| LUAD | CHRNA3 | 1.14 | 13.618 | 1.26e-65 |
| LUAD | ALK | 0.9 | 15.357 | 1.59e-39 |
| LUAD | TP53 | 0.845 | 14.823 | 1.34e-39 |
| UCEC | FBXW7 | -0.878 | 15.289 | 2.31e-17 |

Supplementary Table 3: Confusion matrix for the SVM model on the test set in multi-cancer classification mode (proteome dataset).

| Truth | Normal.CCRCC | Tumor.CCRCC | Normal.COAD | Tumor.COAD | Normal.HNSCC | Tumor.HNSCC | Normal.LSCC | Tumor.LSCC | Normal.LUAD | Tumor.LUAD | Normal.OV | Tumor.OV | Normal.PDAC | Tumor.PDAC | Normal.UCEC | Tumor.UCEC |
| --- | --- | --- | --- | --- | --- | --- | --- | --- | --- | --- | --- | --- | --- | --- | --- | --- |
| Normal.CCRCC | 15 | 0 | 0 | 0 | 0 | 0 | 0 | 0 | 0 | 0 | 0 | 0 | 0 | 0 | 0 | 0 |
| Tumor.CCRCC | 0 | 18 | 0 | 0 | 0 | 0 | 0 | 0 | 0 | 0 | 0 | 0 | 0 | 0 | 0 | 0 |
| Normal.COAD | 0 | 0 | 19 | 0 | 0 | 0 | 0 | 0 | 0 | 0 | 0 | 0 | 0 | 0 | 0 | 0 |
| Tumor.COAD | 0 | 0 | 0 | 16 | 0 | 0 | 0 | 0 | 0 | 0 | 0 | 0 | 0 | 0 | 0 | 0 |
| Normal.HNSCC | 0 | 0 | 0 | 0 | 11 | 0 | 0 | 0 | 0 | 0 | 0 | 0 | 0 | 0 | 0 | 0 |
| Tumor.HNSCC | 0 | 0 | 0 | 0 | 0 | 19 | 0 | 0 | 0 | 0 | 0 | 0 | 0 | 0 | 0 | 0 |
| Normal.LSCC | 0 | 0 | 0 | 0 | 0 | 0 | 19 | 0 | 0 | 0 | 0 | 0 | 0 | 0 | 0 | 0 |
| Tumor.LSCC | 0 | 0 | 0 | 0 | 0 | 0 | 0 | 22 | 0 | 0 | 0 | 0 | 0 | 0 | 0 | 0 |
| Normal.LUAD | 0 | 0 | 0 | 0 | 0 | 0 | 0 | 0 | 17 | 0 | 0 | 0 | 0 | 0 | 0 | 0 |
| Tumor.LUAD | 0 | 0 | 0 | 0 | 0 | 0 | 0 | 0 | 0 | 18 | 0 | 0 | 0 | 0 | 0 | 0 |
| Normal.OV | 0 | 0 | 0 | 0 | 0 | 0 | 0 | 0 | 0 | 0 | 3 | 0 | 0 | 0 | 0 | 0 |
| Tumor.OV | 0 | 0 | 0 | 0 | 0 | 0 | 0 | 0 | 0 | 0 | 0 | 16 | 0 | 0 | 0 | 0 |
| Normal.PDAC | 0 | 0 | 0 | 0 | 0 | 0 | 0 | 0 | 0 | 0 | 0 | 0 | 9 | 0 | 0 | 0 |
| Tumor.PDAC | 0 | 0 | 0 | 0 | 0 | 0 | 0 | 0 | 0 | 0 | 0 | 0 | 0 | 20 | 0 | 0 |
| Normal.UCEC | 0 | 0 | 0 | 0 | 0 | 0 | 0 | 0 | 0 | 0 | 0 | 0 | 0 | 0 | 4 | 0 |
| Tumor.UCEC | 0 | 0 | 0 | 0 | 0 | 0 | 0 | 0 | 0 | 0 | 0 | 0 | 0 | 0 | 0 | 17 |

Supplementary Table 4: Confusion matrix for the SVM model on validation sets (PDAC and HNSCC) in multi-cancer classification mode (proteome dataset).

| y_true | Normal.HNSCC | Normal.PDAC | Tumor.HNSCC | Tumor.PDAC |
| --- | --- | --- | --- | --- |
| Normal.HNSCC | 63 | 0 | 0 | 0 |
| Normal.PDAC | 0 | 75 | 0 | 0 |
| Tumor.HNSCC | 0 | 0 | 109 | 0 |
| Tumor.PDAC | 0 | 6 | 0 | 134 |

Supplementary Table 5: Performance metrics of the SVM model on validation sets (PDAC and HNSCC) in multi-cancer classification mode (proteome dataset).

| Class | Accuracy | Precision | Sensitivity | F1_Score | Specificity | AUC_average |
| --- | --- | --- | --- | --- | --- | --- |
| Normal.HNSCC | 1 | 1 | 1 | 1 | 1 | 0.996 |
| Normal.PDAC | 0.984 | 0.926 | 1 | 0.962 | 0.981 | 0.996 |
| Tumor.HNSCC | 1 | 1 | 1 | 1 | 1 | 0.996 |
| Tumor.PDAC | 0.984 | 1 | 0.957 | 0.978 | 1 | 0.996 |

Supplementary Table 6: Confusion matrix for the RF model on the test set in multi-cancer classification mode (proteome dataset).

| Truth | Normal.CCRCC | Tumor.CCRCC | Normal.COAD | Tumor.COAD | Normal.HNSCC | Tumor.HNSCC | Normal.LSCC | Tumor.LSCC | Normal.LUAD | Tumor.LUAD | Normal.OV | Tumor.OV | Normal.PDAC | Tumor.PDAC | Normal.UCEC | Tumor.UCEC |
| --- | --- | --- | --- | --- | --- | --- | --- | --- | --- | --- | --- | --- | --- | --- | --- | --- |
| Normal.CCRCC | 15 | 0 | 0 | 0 | 0 | 0 | 0 | 0 | 0 | 0 | 0 | 0 | 0 | 0 | 0 | 0 |
| Tumor.CCRCC | 0 | 18 | 0 | 0 | 0 | 0 | 0 | 0 | 0 | 0 | 0 | 0 | 0 | 0 | 0 | 0 |
| Normal.COAD | 0 | 0 | 19 | 0 | 0 | 0 | 0 | 0 | 0 | 0 | 0 | 0 | 0 | 0 | 0 | 0 |
| Tumor.COAD | 0 | 0 | 0 | 16 | 0 | 0 | 0 | 0 | 0 | 0 | 0 | 0 | 0 | 0 | 0 | 0 |
| Normal.HNSCC | 0 | 0 | 0 | 0 | 11 | 0 | 0 | 0 | 0 | 0 | 0 | 0 | 0 | 0 | 0 | 0 |
| Tumor.HNSCC | 0 | 0 | 0 | 0 | 0 | 19 | 0 | 0 | 0 | 0 | 0 | 0 | 0 | 0 | 0 | 0 |
| Normal.LSCC | 0 | 0 | 0 | 0 | 0 | 0 | 19 | 0 | 0 | 0 | 0 | 0 | 0 | 0 | 0 | 0 |
| Tumor.LSCC | 0 | 0 | 0 | 0 | 0 | 0 | 0 | 22 | 0 | 0 | 0 | 0 | 0 | 0 | 0 | 0 |
| Normal.LUAD | 0 | 0 | 0 | 0 | 0 | 0 | 0 | 0 | 17 | 0 | 0 | 0 | 0 | 0 | 0 | 0 |
| Tumor.LUAD | 0 | 0 | 0 | 0 | 0 | 0 | 0 | 0 | 0 | 18 | 0 | 0 | 0 | 0 | 0 | 0 |
| Normal.OV | 0 | 0 | 0 | 0 | 0 | 0 | 0 | 0 | 0 | 0 | 3 | 0 | 0 | 0 | 0 | 0 |
| Tumor.OV | 0 | 0 | 0 | 0 | 0 | 0 | 0 | 0 | 0 | 0 | 0 | 16 | 0 | 0 | 0 | 0 |
| Normal.PDAC | 0 | 0 | 0 | 0 | 0 | 0 | 0 | 0 | 0 | 0 | 0 | 0 | 9 | 0 | 0 | 0 |
| Tumor.PDAC | 0 | 0 | 0 | 0 | 0 | 0 | 0 | 0 | 0 | 0 | 0 | 0 | 0 | 20 | 0 | 0 |
| Normal.UCEC | 0 | 0 | 0 | 0 | 0 | 0 | 0 | 0 | 0 | 0 | 0 | 0 | 0 | 0 | 4 | 0 |
| Tumor.UCEC | 0 | 0 | 0 | 0 | 0 | 0 | 0 | 0 | 0 | 0 | 0 | 0 | 0 | 0 | 0 | 17 |

Supplementary Table 7: Performance metrics of the RF model on the test set in multi-cancer classification mode (proteome dataset).

| Class | Accuracy | Precision | Sensitivity | F1_Score | Specificity | AUC_average |
| --- | --- | --- | --- | --- | --- | --- |
| Normal.CCRCC | 1 | 1 | 1 | 1 | 1 | 1 |
| Tumor.CCRCC | 1 | 1 | 1 | 1 | 1 | 1 |
| Normal.COAD | 1 | 1 | 1 | 1 | 1 | 1 |
| Tumor.COAD | 1 | 1 | 1 | 1 | 1 | 1 |
| Normal.HNSCC | 1 | 1 | 1 | 1 | 1 | 1 |
| Tumor.HNSCC | 1 | 1 | 1 | 1 | 1 | 1 |
| Normal.LSCC | 1 | 1 | 1 | 1 | 1 | 1 |
| Tumor.LSCC | 1 | 1 | 1 | 1 | 1 | 1 |
| Normal.LUAD | 1 | 1 | 1 | 1 | 1 | 1 |
| Tumor.LUAD | 1 | 1 | 1 | 1 | 1 | 1 |
| Normal.OV | 1 | 1 | 1 | 1 | 1 | 1 |
| Tumor.OV | 1 | 1 | 1 | 1 | 1 | 1 |
| Normal.PDAC | 1 | 1 | 1 | 1 | 1 | 1 |
| Tumor.PDAC | 1 | 1 | 1 | 1 | 1 | 1 |
| Normal.UCEC | 0.997 | 0.837 | 1 | 0.907 | 0.997 | 1 |
| Tumor.UCEC | 0.997 | 1 | 0.958 | 0.979 | 1 | 1 |

Supplementary Table 8: Confusion matrix for the RF model on validation sets (PDAC and HNSCC) in multi-cancer classification mode (proteome dataset).

| y_true | Normal.HNSCC | Normal.PDAC | Tumor.HNSCC | Tumor.PDAC |
| --- | --- | --- | --- | --- |
| Normal.HNSCC | 63 | 0 | 0 | 0 |
| Normal.PDAC | 0 | 73 | 0 | 2 |
| Tumor.HNSCC | 3 | 0 | 106 | 0 |
| Tumor.PDAC | 0 | 4 | 0 | 136 |

Supplementary Table 9: Performance metrics of the RF model on validation sets (PDAC and HNSCC) in multi-cancer classification mode (proteome dataset).

| Class | Accuracy | Precision | Sensitivity | F1_Score | Specificity | AUC_average |
| --- | --- | --- | --- | --- | --- | --- |
| Normal.HNSCC | 0.992 | 0.955 | 1 | 0.977 | 0.991 | 0.993 |
| Normal.PDAC | 0.984 | 0.948 | 0.973 | 0.961 | 0.987 | 0.993 |
| Tumor.HNSCC | 0.992 | 1 | 0.972 | 0.986 | 1 | 0.993 |
| Tumor.PDAC | 0.984 | 0.986 | 0.971 | 0.978 | 0.992 | 0.993 |

Supplementary Table 10: Confusion matrix for the ANN model on the test set in multi-cancer classification mode (proteome dataset).

| Truth | Normal.CCRCC | Tumor.CCRCC | Normal.COAD | Tumor.COAD | Normal.HNSCC | Tumor.HNSCC | Normal.LSCC | Tumor.LSCC | Normal.LUAD | Tumor.LUAD | Normal.OV | Tumor.OV | Normal.PDAC | Tumor.PDAC | Normal.UCEC | Tumor.UCEC |
| --- | --- | --- | --- | --- | --- | --- | --- | --- | --- | --- | --- | --- | --- | --- | --- | --- |
| Normal.CCRCC | 6 | 1 | 0 | 0 | 0 | 0 | 0 | 0 | 0 | 0 | 0 | 0 | 0 | 0 | 0 | 0 |
| Tumor.CCRCC | 0 | 10 | 0 | 0 | 0 | 0 | 0 | 0 | 0 | 0 | 0 | 0 | 0 | 0 | 0 | 0 |
| Normal.COAD | 0 | 0 | 10 | 0 | 0 | 0 | 0 | 0 | 0 | 0 | 0 | 0 | 0 | 0 | 0 | 0 |
| Tumor.COAD | 0 | 0 | 0 | 8 | 0 | 0 | 0 | 0 | 0 | 0 | 0 | 0 | 0 | 0 | 0 | 0 |
| Normal.HNSCC | 0 | 0 | 0 | 0 | 1 | 0 | 0 | 0 | 0 | 0 | 0 | 0 | 0 | 0 | 0 | 0 |
| Tumor.HNSCC | 0 | 0 | 0 | 0 | 0 | 8 | 0 | 0 | 0 | 0 | 0 | 0 | 0 | 0 | 0 | 0 |
| Normal.LSCC | 0 | 0 | 0 | 0 | 0 | 0 | 13 | 0 | 0 | 0 | 0 | 0 | 0 | 0 | 0 | 0 |
| Tumor.LSCC | 0 | 0 | 0 | 0 | 0 | 0 | 0 | 13 | 0 | 0 | 0 | 0 | 0 | 0 | 0 | 0 |
| Normal.LUAD | 0 | 0 | 0 | 0 | 0 | 0 | 0 | 0 | 11 | 0 | 0 | 0 | 0 | 0 | 0 | 0 |
| Tumor.LUAD | 0 | 0 | 0 | 0 | 0 | 0 | 0 | 0 | 0 | 10 | 0 | 0 | 0 | 0 | 0 | 0 |
| Normal.OV | 0 | 0 | 0 | 0 | 0 | 0 | 0 | 0 | 0 | 0 | 0 | 0 | 0 | 0 | 0 | 0 |
| Tumor.OV | 0 | 0 | 0 | 0 | 0 | 0 | 0 | 0 | 0 | 0 | 0 | 11 | 0 | 0 | 0 | 0 |
| Normal.PDAC | 0 | 0 | 0 | 0 | 0 | 0 | 0 | 0 | 0 | 0 | 0 | 0 | 4 | 0 | 0 | 0 |
| Tumor.PDAC | 0 | 0 | 0 | 0 | 0 | 0 | 0 | 0 | 0 | 0 | 0 | 0 | 0 | 9 | 0 | 0 |
| Normal.UCEC | 0 | 0 | 0 | 0 | 0 | 0 | 0 | 0 | 0 | 0 | 0 | 0 | 0 | 0 | 2 | 0 |
| Tumor.UCEC | 0 | 0 | 0 | 0 | 0 | 0 | 0 | 0 | 0 | 0 | 0 | 0 | 0 | 0 | 0 | 5 |

Supplementary Table 11: Performance metrics of the ANN model on the test set in multi-cancer classification mode (proteome dataset).

| Class | Accuracy | Precision | Sensitivity | F1_Score | Specificity | AUC_average |
| --- | --- | --- | --- | --- | --- | --- |
| Normal.CCRCC | 0.992 | 0.857 | 1 | 0.923 | 0.991 | 1 |
| Tumor.CCRCC | 0.992 | 1 | 0.909 | 0.952 | 1 | 1 |
| Normal.COAD | 1 | 1 | 1 | 1 | 1 | 1 |
| Tumor.COAD | 1 | 1 | 1 | 1 | 1 | 1 |
| Normal.HNSCC | 1 | 1 | 1 | 1 | 1 | 1 |
| Tumor.HNSCC | 1 | 1 | 1 | 1 | 1 | 1 |
| Normal.LSCC | 1 | 1 | 1 | 1 | 1 | 1 |
| Tumor.LSCC | 1 | 1 | 1 | 1 | 1 | 1 |
| Normal.LUAD | 1 | 1 | 1 | 1 | 1 | 1 |
| Tumor.LUAD | 1 | 1 | 1 | 1 | 1 | 1 |
| Tumor.OV | 1 | 1 | 1 | 1 | 1 | 1 |
| Normal.PDAC | 1 | 1 | 1 | 1 | 1 | 1 |
| Tumor.PDAC | 1 | 1 | 1 | 1 | 1 | 1 |
| Normal.UCEC | 1 | 1 | 1 | 1 | 1 | 1 |
| Tumor.UCEC | 1 | 1 | 1 | 1 | 1 | 1 |

Supplementary Table 12: Confusion matrix for the ANN model on validation sets (PDAC and HNSCC) in multi-cancer classification mode (proteome dataset).

| y_true | Normal.HNSCC | Normal.PDAC | Tumor.HNSCC | Tumor.PDAC |
| --- | --- | --- | --- | --- |
| Normal.HNSCC | 63 | 0 | 0 | 0 |
| Normal.PDAC | 0 | 73 | 0 | 2 |
| Tumor.HNSCC | 3 | 0 | 106 | 0 |
| Tumor.PDAC | 0 | 4 | 0 | 136 |

Supplementary Table 13: Performance metrics of the ANN model on validation sets (PDAC and HNSCC) in multi-cancer classification mode (proteome dataset).

| Class | Accuracy | Precision | Sensitivity | F1_Score | Specificity | AUC_average |
| --- | --- | --- | --- | --- | --- | --- |
| Normal.HNSCC | 0.992 | 0.955 | 1 | 0.977 | 0.991 | 0.993 |
| Normal.PDAC | 0.984 | 0.948 | 0.973 | 0.961 | 0.987 | 0.993 |
| Tumor.HNSCC | 0.992 | 1 | 0.972 | 0.986 | 1 | 0.993 |
| Tumor.PDAC | 0.984 | 0.986 | 0.971 | 0.978 | 0.992 | 0.993 |

Supplementary Table 14: Confusion matrix for all three models on the HNSCC dataset as a validation set in single-cancer classification mode (proteome dataset).

| y_true | Normal.HNSCC | Tumor.HNSCC | Model |
| --- | --- | --- | --- |
| Normal.HNSCC | 63 | 0 | SVM |
| Tumor.HNSCC | 1 | 108 |  |
| Normal.HNSCC | 63 | 0 | RF |
| Tumor.HNSCC | 1 | 108 |  |
| Normal.HNSCC | 63 | 0 | ANN |
| Tumor.HNSCC | 1 | 108 |  |

Supplementary Table 15: Performance metrics of all three models on the HNSCC dataset as a validation set in single-cancer classification mode (proteome dataset).

| Accuracy | Precision | Sensitivity | F1_Score | Specificity | AUC | Model |
| --- | --- | --- | --- | --- | --- | --- |
| 0.994 | 1 | 0.984 | 0.992 | 1 | 0.995 | SVM |
| 0.994 | 1 | 0.984 | 0.992 | 1 | 0.995 | RF |
| 0.994 | 1 | 0.984 | 0.992 | 1 | 0.995 | ANN |

Supplementary Table 16: Confusion matrix for all three models on the PDAC dataset as a validation set in single-cancer classification mode (proteome dataset).

| y_true | Normal.PDAC | Tumor.PDAC | Model |
| --- | --- | --- | --- |
| Normal.PDAC | 72 | 3 | SVM |
| Tumor.PDAC | 3 | 137 |  |
| Normal.PDAC | 72 | 3 | RF |
| Tumor.PDAC | 3 | 137 |  |
| Normal.PDAC | 72 | 3 | ANN |
| Tumor.PDAC | 3 | 137 |  |

Supplementary Table 17: Performance metrics of all three models on the PDAC dataset as a validation set in single-cancer classification mode (proteome dataset).

| Accuracy | Precision | Sensitivity | F1_Score | Specificity | AUC | Model |
| --- | --- | --- | --- | --- | --- | --- |
| 0.972 | 0.96 | 0.96 | 0.96 | 0.979 | 0.969 | SVM |
| 0.972 | 0.96 | 0.96 | 0.96 | 0.979 | 0.969 | RF |
| 0.972 | 0.96 | 0.96 | 0.96 | 0.979 | 0.969 | ANN |

Supplementary Table 18: Confusion matrix of the SVM model on the test set for phosphoproteome dataset (multi-cancer mode).

| Truth | Normal.CCRCC | Tumor.CCRCC | Normal.COAD | Tumor.COAD | Normal.HNSCC | Tumor.HNSCC | Normal.LSCC | Tumor.LSCC | Normal.LUAD | Tumor.LUAD | Normal.OV | Tumor.OV | Normal.PDAC | Tumor.PDAC | Normal.UCEC | Tumor.UCEC |
| --- | --- | --- | --- | --- | --- | --- | --- | --- | --- | --- | --- | --- | --- | --- | --- | --- |
| Normal.CCRCC | 15 | 0 | 0 | 0 | 0 | 0 | 0 | 0 | 0 | 0 | 0 | 0 | 0 | 0 | 0 | 0 |
| Tumor.CCRCC | 0 | 20 | 0 | 0 | 0 | 0 | 0 | 0 | 0 | 0 | 0 | 0 | 0 | 0 | 0 | 0 |
| Normal.COAD | 0 | 0 | 16 | 0 | 0 | 0 | 0 | 0 | 0 | 0 | 0 | 0 | 0 | 0 | 0 | 0 |
| Tumor.COAD | 0 | 0 | 0 | 18 | 0 | 0 | 0 | 0 | 0 | 0 | 0 | 0 | 0 | 0 | 0 | 0 |
| Normal.HNSCC | 0 | 0 | 0 | 0 | 11 | 0 | 0 | 0 | 0 | 0 | 0 | 0 | 0 | 0 | 0 | 0 |
| Tumor.HNSCC | 0 | 0 | 0 | 0 | 0 | 18 | 0 | 0 | 0 | 0 | 0 | 0 | 0 | 0 | 0 | 0 |
| Normal.LSCC | 0 | 0 | 0 | 0 | 0 | 0 | 19 | 0 | 0 | 0 | 0 | 0 | 0 | 0 | 0 | 0 |
| Tumor.LSCC | 0 | 0 | 0 | 0 | 0 | 0 | 0 | 18 | 0 | 0 | 0 | 0 | 0 | 0 | 0 | 0 |
| Normal.LUAD | 0 | 0 | 0 | 0 | 0 | 0 | 0 | 0 | 17 | 0 | 0 | 0 | 0 | 0 | 0 | 0 |
| Tumor.LUAD | 0 | 0 | 0 | 0 | 0 | 0 | 0 | 0 | 0 | 17 | 0 | 0 | 0 | 0 | 0 | 0 |
| Normal.OV | 0 | 0 | 0 | 0 | 0 | 0 | 0 | 0 | 0 | 0 | 2 | 0 | 0 | 0 | 0 | 0 |
| Tumor.OV | 0 | 0 | 0 | 0 | 0 | 0 | 0 | 0 | 0 | 0 | 0 | 16 | 0 | 0 | 0 | 0 |
| Normal.PDAC | 0 | 0 | 0 | 0 | 0 | 0 | 0 | 0 | 0 | 0 | 0 | 0 | 9 | 0 | 0 | 0 |
| Tumor.PDAC | 0 | 0 | 0 | 0 | 0 | 0 | 0 | 0 | 0 | 0 | 0 | 0 | 0 | 19 | 0 | 0 |
| Normal.UCEC | 0 | 0 | 0 | 0 | 0 | 0 | 0 | 0 | 0 | 0 | 0 | 0 | 0 | 0 | 4 | 0 |
| Tumor.UCEC | 0 | 0 | 0 | 0 | 0 | 0 | 0 | 0 | 0 | 0 | 0 | 0 | 0 | 0 | 0 | 16 |

Supplementary Table 19: Performance metrics of the SVM model on the test set for phosphoproteome dataset (multi-cancer mode).

| Class | Accuracy | Precision | Sensitivity | F1_Score | Specificity | AUC_average |
| --- | --- | --- | --- | --- | --- | --- |
| Normal.CCRCC | 1 | 1 | 1 | 1 | 1 | 1 |
| Tumor.CCRCC | 1 | 1 | 1 | 1 | 1 | 1 |
| Normal.COAD | 1 | 1 | 0.995 | 0.997 | 1 | 1 |
| Tumor.COAD | 1 | 0.995 | 1 | 0.997 | 1 | 1 |
| Normal.HNSCC | 1 | 1 | 1 | 1 | 1 | 1 |
| Tumor.HNSCC | 1 | 1 | 1 | 1 | 1 | 1 |
| Normal.LSCC | 1 | 1 | 1 | 1 | 1 | 1 |
| Tumor.LSCC | 1 | 1 | 1 | 1 | 1 | 1 |
| Normal.LUAD | 1 | 1 | 1 | 1 | 1 | 1 |
| Tumor.LUAD | 1 | 1 | 1 | 1 | 1 | 1 |
| Normal.OV | 1 | 1 | 1 | 1 | 1 | 1 |
| Tumor.OV | 1 | 1 | 1 | 1 | 1 | 1 |
| Normal.PDAC | 1 | 1 | 1 | 1 | 1 | 1 |
| Tumor.PDAC | 1 | 1 | 1 | 1 | 1 | 1 |
| Normal.UCEC | 1 | 1 | 1 | 1 | 1 | 1 |
| Tumor.UCEC | 1 | 1 | 1 | 1 | 1 | 1 |

Supplementary Table 20: Confusion matrix of the RF model on the test set for phosphoproteome dataset (multi-cancer mode).

| Truth | Normal.CCRCC | Tumor.CCRCC | Normal.COAD | Tumor.COAD | Normal.HNSCC | Tumor.HNSCC | Normal.LSCC | Tumor.LSCC | Normal.LUAD | Tumor.LUAD | Normal.OV | Tumor.OV | Normal.PDAC | Tumor.PDAC | Normal.UCEC | Tumor.UCEC |
| --- | --- | --- | --- | --- | --- | --- | --- | --- | --- | --- | --- | --- | --- | --- | --- | --- |
| Normal.CCRCC | 15 | 0 | 0 | 0 | 0 | 0 | 0 | 0 | 0 | 0 | 0 | 0 | 0 | 0 | 0 | 0 |
| Tumor.CCRCC | 0 | 20 | 0 | 0 | 0 | 0 | 0 | 0 | 0 | 0 | 0 | 0 | 0 | 0 | 0 | 0 |
| Normal.COAD | 0 | 0 | 15 | 0 | 0 | 0 | 0 | 0 | 0 | 0 | 0 | 0 | 0 | 0 | 0 | 0 |
| Tumor.COAD | 0 | 0 | 1 | 18 | 0 | 0 | 0 | 0 | 0 | 0 | 0 | 0 | 0 | 0 | 0 | 0 |
| Normal.HNSCC | 0 | 0 | 0 | 0 | 11 | 0 | 0 | 0 | 0 | 0 | 0 | 0 | 0 | 0 | 0 | 0 |
| Tumor.HNSCC | 0 | 0 | 0 | 0 | 0 | 18 | 0 | 0 | 0 | 0 | 0 | 0 | 0 | 0 | 0 | 0 |
| Normal.LSCC | 0 | 0 | 0 | 0 | 0 | 0 | 19 | 0 | 0 | 0 | 0 | 0 | 0 | 0 | 0 | 0 |
| Tumor.LSCC | 0 | 0 | 0 | 0 | 0 | 0 | 0 | 18 | 0 | 0 | 0 | 0 | 0 | 0 | 0 | 0 |
| Normal.LUAD | 0 | 0 | 0 | 0 | 0 | 0 | 0 | 0 | 17 | 0 | 0 | 0 | 0 | 0 | 0 | 0 |
| Tumor.LUAD | 0 | 0 | 0 | 0 | 0 | 0 | 0 | 0 | 0 | 17 | 0 | 0 | 0 | 0 | 0 | 0 |
| Normal.OV | 0 | 0 | 0 | 0 | 0 | 0 | 0 | 0 | 0 | 0 | 2 | 0 | 0 | 0 | 0 | 0 |
| Tumor.OV | 0 | 0 | 0 | 0 | 0 | 0 | 0 | 0 | 0 | 0 | 0 | 16 | 0 | 0 | 0 | 0 |
| Normal.PDAC | 0 | 0 | 0 | 0 | 0 | 0 | 0 | 0 | 0 | 0 | 0 | 0 | 9 | 0 | 0 | 0 |
| Tumor.PDAC | 0 | 0 | 0 | 0 | 0 | 0 | 0 | 0 | 0 | 0 | 0 | 0 | 0 | 19 | 0 | 0 |
| Normal.UCEC | 0 | 0 | 0 | 0 | 0 | 0 | 0 | 0 | 0 | 0 | 0 | 0 | 0 | 0 | 4 | 0 |
| Tumor.UCEC | 0 | 0 | 0 | 0 | 0 | 0 | 0 | 0 | 0 | 0 | 0 | 0 | 0 | 0 | 0 | 16 |

Supplementary Table 21: Performance metrics of the RF model on the test set for phosphoproteome dataset (multi-cancer mode).

| Class | Accuracy | Precision | Sensitivity | F1_Score | Specificity | AUC_average |
| --- | --- | --- | --- | --- | --- | --- |
| Normal.CCRCC | 1 | 0.994 | 1 | 0.997 | 1 | 1 |
| Tumor.CCRCC | 1 | 1 | 0.995 | 0.997 | 1 | 1 |
| Normal.COAD | 0.994 | 0.995 | 0.921 | 0.956 | 1 | 1 |
| Tumor.COAD | 0.994 | 0.925 | 0.994 | 0.958 | 0.994 | 1 |
| Normal.HNSCC | 1 | 1 | 1 | 1 | 1 | 1 |
| Tumor.HNSCC | 1 | 1 | 1 | 1 | 1 | 1 |
| Normal.LSCC | 1 | 1 | 1 | 1 | 1 | 1 |
| Tumor.LSCC | 1 | 1 | 1 | 1 | 1 | 1 |
| Normal.LUAD | 1 | 1 | 1 | 1 | 1 | 1 |
| Tumor.LUAD | 1 | 1 | 1 | 1 | 1 | 1 |
| Normal.OV | 1 | 1 | 1 | 1 | 1 | 1 |
| Tumor.OV | 1 | 1 | 1 | 1 | 1 | 1 |
| Normal.PDAC | 1 | 1 | 1 | 1 | 1 | 1 |
| Tumor.PDAC | 1 | 1 | 1 | 1 | 1 | 1 |
| Normal.UCEC | 1 | 1 | 1 | 1 | 1 | 1 |
| Tumor.UCEC | 1 | 1 | 1 | 1 | 1 | 1 |

Supplementary Table 22: Confusion matrix of the ANN model on the test set for phosphoproteome dataset (multi-cancer mode).

| Truth | Normal.CCRCC | Tumor.CCRCC | Normal.COAD | Tumor.COAD | Normal.HNSCC | Tumor.HNSCC | Normal.LSCC | Tumor.LSCC | Normal.LUAD | Tumor.LUAD | Normal.OV | Tumor.OV | Normal.PDAC | Tumor.PDAC | Normal.UCEC | Tumor.UCEC |
| --- | --- | --- | --- | --- | --- | --- | --- | --- | --- | --- | --- | --- | --- | --- | --- | --- |
| Normal.CCRCC | 6 | 0 | 0 | 0 | 0 | 0 | 0 | 0 | 0 | 0 | 0 | 0 | 0 | 0 | 0 | 0 |
| Tumor.CCRCC | 0 | 11 | 0 | 0 | 0 | 0 | 0 | 0 | 0 | 0 | 0 | 0 | 0 | 0 | 0 | 0 |
| Normal.COAD | 0 | 0 | 10 | 0 | 0 | 0 | 0 | 0 | 0 | 0 | 0 | 0 | 0 | 0 | 0 | 0 |
| Tumor.COAD | 0 | 0 | 0 | 8 | 0 | 0 | 0 | 0 | 0 | 0 | 0 | 0 | 0 | 0 | 0 | 0 |
| Normal.HNSCC | 0 | 0 | 0 | 0 | 1 | 0 | 0 | 0 | 0 | 0 | 0 | 0 | 0 | 0 | 0 | 0 |
| Tumor.HNSCC | 0 | 0 | 0 | 0 | 0 | 8 | 0 | 0 | 0 | 0 | 0 | 0 | 0 | 0 | 0 | 0 |
| Normal.LSCC | 0 | 0 | 0 | 0 | 0 | 0 | 13 | 0 | 0 | 0 | 0 | 0 | 0 | 0 | 0 | 0 |
| Tumor.LSCC | 0 | 0 | 0 | 0 | 0 | 0 | 0 | 12 | 0 | 0 | 0 | 0 | 0 | 0 | 0 | 0 |
| Normal.LUAD | 0 | 0 | 0 | 0 | 0 | 0 | 0 | 0 | 8 | 0 | 0 | 0 | 0 | 0 | 0 | 0 |
| Tumor.LUAD | 0 | 0 | 0 | 0 | 0 | 0 | 0 | 0 | 0 | 9 | 0 | 0 | 0 | 0 | 0 | 0 |
| Normal.OV | 0 | 0 | 0 | 0 | 0 | 0 | 0 | 0 | 0 | 0 | 3 | 0 | 0 | 0 | 0 | 0 |
| Tumor.OV | 0 | 0 | 0 | 0 | 0 | 0 | 0 | 0 | 0 | 0 | 0 | 13 | 0 | 0 | 0 | 0 |
| Normal.PDAC | 0 | 0 | 0 | 0 | 0 | 0 | 0 | 0 | 0 | 0 | 0 | 0 | 2 | 0 | 0 | 0 |
| Tumor.PDAC | 0 | 0 | 0 | 0 | 0 | 0 | 0 | 0 | 0 | 0 | 0 | 0 | 1 | 7 | 0 | 0 |
| Normal.UCEC | 0 | 0 | 0 | 0 | 0 | 0 | 0 | 0 | 0 | 0 | 0 | 0 | 0 | 0 | 0 | 0 |
| Tumor.UCEC | 0 | 0 | 0 | 0 | 0 | 0 | 0 | 0 | 0 | 0 | 0 | 0 | 0 | 0 | 0 | 8 |

Supplementary Table 23: Performance metrics of the ANN model on the test set for phosphoproteome dataset (multi-cancer mode).

| Class | Accuracy | Precision | Sensitivity | F1_Score | Specificity | AUC_average |
| --- | --- | --- | --- | --- | --- | --- |
| Normal.CCRCC | 1 | 1 | 1 | 1 | 1 | 0.998 |
| Tumor.CCRCC | 1 | 1 | 1 | 1 | 1 | 0.998 |
| Normal.COAD | 1 | 1 | 1 | 1 | 1 | 0.998 |
| Tumor.COAD | 1 | 1 | 1 | 1 | 1 | 0.998 |
| Normal.HNSCC | 1 | 1 | 1 | 1 | 1 | 0.998 |
| Tumor.HNSCC | 1 | 1 | 1 | 1 | 1 | 0.998 |
| Normal.LSCC | 1 | 1 | 1 | 1 | 1 | 0.998 |
| Tumor.LSCC | 1 | 1 | 1 | 1 | 1 | 0.998 |
| Normal.LUAD | 1 | 1 | 1 | 1 | 1 | 0.998 |
| Tumor.LUAD | 1 | 1 | 1 | 1 | 1 | 0.998 |
| Normal.OV | 1 | 1 | 1 | 1 | 1 | 0.998 |
| Tumor.OV | 1 | 1 | 1 | 1 | 1 | 0.998 |
| Normal.PDAC | 0.992 | 1 | 0.667 | 0.8 | 1 | 0.998 |
| Tumor.PDAC | 0.992 | 0.875 | 1 | 0.933 | 0.991 | 0.998 |
| Tumor.UCEC | 1 | 1 | 1 | 1 | 1 | 0.998 |

Supplementary Table 24: Confusion matrices of SVM, RF, and ANN models in single-cancer mode (CCRCC) on the phosphoproteome dataset.

| y_true | Normal.CCRCC | Tumor.CCRCC | Model |
| --- | --- | --- | --- |
| Normal.CCRCC | 15 | 0 | SVM |
| Tumor.CCRCC | 0 | 18 |  |
| Normal.CCRCC | 15 | 0 | RF |
| Tumor.CCRCC | 0 | 18 |  |
| Normal.CCRCC | 6 | 0 | ANN |
| Tumor.CCRCC | 0 | 11 |  |

Supplementary Table 25: Performance metrics of SVM, RF, and ANN models in single-cancer mode (CCRCC) on the phosphoproteome dataset.

| Accuracy | Precision | Sensitivity | F1_Score | Specificity | AUC |  |
| --- | --- | --- | --- | --- | --- | --- |
| 1 | 1 | 1 | 1 | 1 | 1 | SVM |
| 1 | 1 | 1 | 1 | 1 | 1 | RF |
| 1 | 1 | 1 | 1 | 1 | 1 | ANN |
